# Evaluating the consequences of common assumptions in run reconstructions on Pacific-salmon biological status assessments

**DOI:** 10.1101/868927

**Authors:** Stephanie J. Peacock, Eric Hertz, Carrie A. Holt, Brendan Connors, Cameron Freshwater, Katrina Connors

## Abstract

Information on biological status is essential for designing, implementing, and evaluating management strategies and recovery plans for threatened or exploited species. However, the data required to quantify status are often limited, and it is important to understand how assessments of status may be biased by assumptions in data analysis. For Pacific salmon, biological status assessments based on spawner abundances and spawner-recruitment (SR) analyses often involve “run reconstructions” that impute missing spawner data, expand observed spawner abundance to account for unmonitored streams, assign catch to individual stocks, and quantify age-at-return. Using a stochastic simulation approach, we quantified how common assumptions in run reconstructions biased assessments of biological status based on spawner abundance. We found that status assessments were robust to most common assumptions in run reconstructions, even in the face of declining monitoring coverage, but that overestimating catch tended to increase rates of status misclassification. Our results lend confidence to biological status assessments based on spawner abundances and SR analyses, even in the face of incomplete data.

## Introduction

Timely and effective management of fish and wildlife relies on accurate information about the current biological status of populations. Status assessments form the basis of many conservation decision-support tools, though they have limited influence without clear and binding linkages to policy and decision making. Methods to determine the allocation of resources that maximize biodiversity conservation, such as Priority Threat Management (Martin et al. 2018; Carwardine et al. 2019; Walsh et al. 2020), are most effective when baseline information on status can help identify which threatened populations are at highest risk of extirpation without management intervention, and which are most likely to benefit from recovery planning given limited resources. Metrics of biological status may also be useful for defining recovery goals and as performance measures when evaluating the effectiveness of different management strategies (e.g., Walsh et al. 2020). Knowledge of current biological status is therefore needed for both implementing and evaluating management strategies and recovery plans.

Despite the recognized need for information on biological status, complete data on abundance, trends, and demographic parameters are rarely available for populations or species that need to be assessed (IUCN 2017). Improving the analytical approaches used to quantify status is one way to reduce assumptions and limit the bias in status outcomes from low quality or quantity data (e.g., Fleischman et al. 2013; Staton et al. 2020), but the methods widely applied in assessing fish and wildlife can be slow to change (Peterman 2018) and analytical resources are sometimes not available to implement new advanced approaches on all populations or species of interest. Thus, evaluating the impact of assumptions in the approaches that are widely applied is critical to understanding the potential biases, uncertainty, and limitations of status assessments (Chen et al. 2003; Wetzel and Punt 2011).

Closed-loop simulation models are useful tools for evaluating the potential biases in status assessments due to various ecological and management uncertainties (Walters 1986). The major advantages of simulation approaches (over, for example, retrospective analyses) are that the model can be tailored to different biological and management scenarios, allowing management strategy evaluation (e.g., Punt 1992; Holt and Peterman 2006, 2008), and the true status of the system is known perfectly, allowing quantification of bias in estimated parameters (e.g., Pyper and Peterman 1998; Dorner et al. 2009) and estimated status (e.g., Peacock and Holt 2012; Holt and Folkes 2015; Holt et al. 2018). Simulation approaches are extremely flexible and enable testing common assumptions that underpin status assessments at broad spatial scales in the face imperfect information, and have been applied extensively in fisheries research. In this study, we apply a simulation approach to understanding potential biases in biological status assessments of Pacific salmon introduced by assumptions in the analysis of spawner and recruitment data.

### Assessing biological status of Pacific salmon

Pacific salmon (*Oncorhynchus* spp.) are a highly exploited group of species and many populations have experienced declines in recent decades due to overfishing, changing ocean conditions, and freshwater habitat degradation (e.g., COSEWIC 2016, 2017; Brown et al. 2019). Pacific salmon are anadromous and semelparous, returning from ocean rearing grounds to spawn in freshwater before dying, and are typically vulnerable to fisheries upon their return to coastal waters. The data required to assess biological status of Pacific salmon can include annual estimates of the number of returning adult salmon to individual rivers, fisheries catch or harvest rates, and the age composition of returning salmon. Often, these data are incomplete and require imputation. “Run reconstructions” have been undertaken for many salmon stocks to expand spawner abundances to account for unmonitored streams and estimate the abundance of recruits to coastal fisheries prior to spawning (i.e., recruitment; Cave and Gazey 1994; English et al. 2007, 2016, 2018). The exact procedure undertaken depends on the life-history traits and available data for each population, and can include complexities such as spatial and temporal variability in returns among spawning populations. Status assessments that rely on run reconstructions have been adopted by, for example, local management organisations, the Marine Stewardship Council (www.msc.org), the Pacific Salmon Treaty (PSC 2019), COSEWIC (e.g., COSEWIC 2016, 2017), and the Pacific Salmon Foundation (Connors et al. 2013, 2018, 2019).

Fisheries and Oceans Canada (DFO) also relies on run reconstructions when assessing the biological status of Conservation Units (CUs) – groups of wild salmon that, if lost, are unlikely to recolonize within an acceptable timeframe – under Canada’s Wild Salmon Policy (WSP; Fisheries and Oceans Canada 2005). Under the WSP’s biological status assessment framework, quantifiable metrics are calculated from available data and compared against biological reference points, or “benchmarks”, to arrive at a status outcome of red, amber, or green (Fisheries and Oceans Canada 2005; Figure 1). A red status indicates that a CU has low spawner abundance and/or reduced spatial distribution and management intervention is required to avoid extirpation. A green status indicates that the CU is able to sustain maximum annual catch under existing environmental conditions. The specific benchmarks delineating these status zones must consider uncertainties in metrics and the unique biological characteristics (e.g., age composition) of the CU being assessed. Indeed, in their “integrated status assessments”, DFO engages experts to consider the unique context of each CU and the quality and quantity of data that may be used to estimate status (e.g., DFO 2015, 2016, 2018a). However, the required resources and time (typically 1-3 years) have meant that integrated status assessments have been completed for only 9% of CUs since the WSP was released nearly 15 years ago, and reports are often 2-4 years out of date when they are released (DFO 2019).

**Figure 1.**
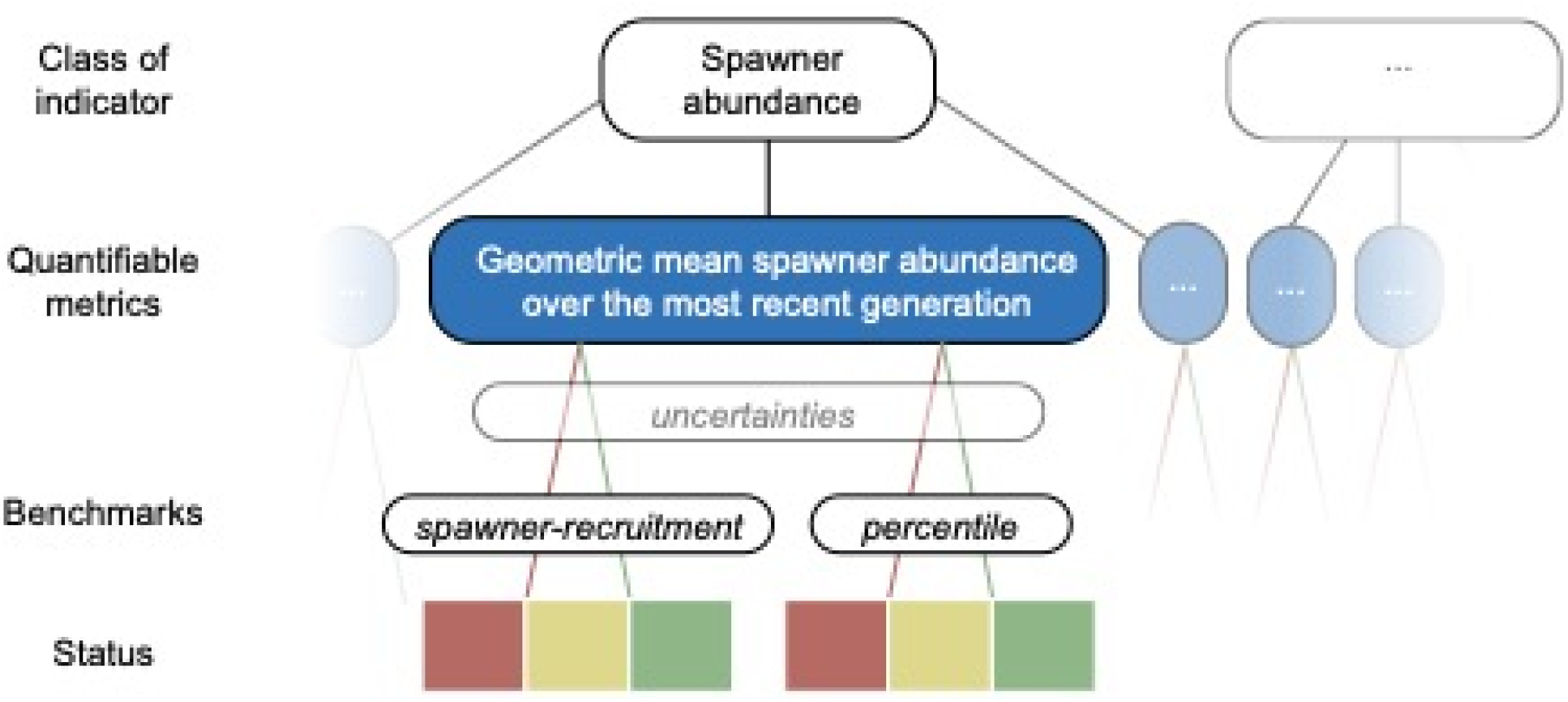
Illustration of the WSP status assessment framework (adapted from Holt et al. 2009). We focused on the geometric mean spawner abundance (metric, blue) under the spawner abundance indicator. This metric was assessed against two types of benchmarks: spawner-recruitment (SR) and percentile (see Figure 2). Faded boxes represent other types of metrics and indicators that may be included in integrated status assessments but were beyond the scope of what we considered.

Data-driven approaches focused on calculating quantitative metrics without expert elicitation can provide timely status updates that are standardized and comparable across CUs using transparent and repeatable methodology. For example, the Pacific Salmon Foundation (PSF) has undertaken a widespread effort to apply a data-driven approach to assessing spawner abundance under the WSP framework, with results for the north and central coast openly available through their Pacific Salmon Explorer (PSE) – www.salmonexplorer.ca (Connors et al. 2013, 2018, 2019), and are currently expanding their assessments to all remaining salmon CUs in British Columbia (BC). Despite the benefits of this approach, multiple assumptions in run reconstructions (Table 1) mean that potential biases can compound, and the impact on status outcomes needs to be quantified to lend confidence and credibility to data-driven assessments.

**Table 1.**
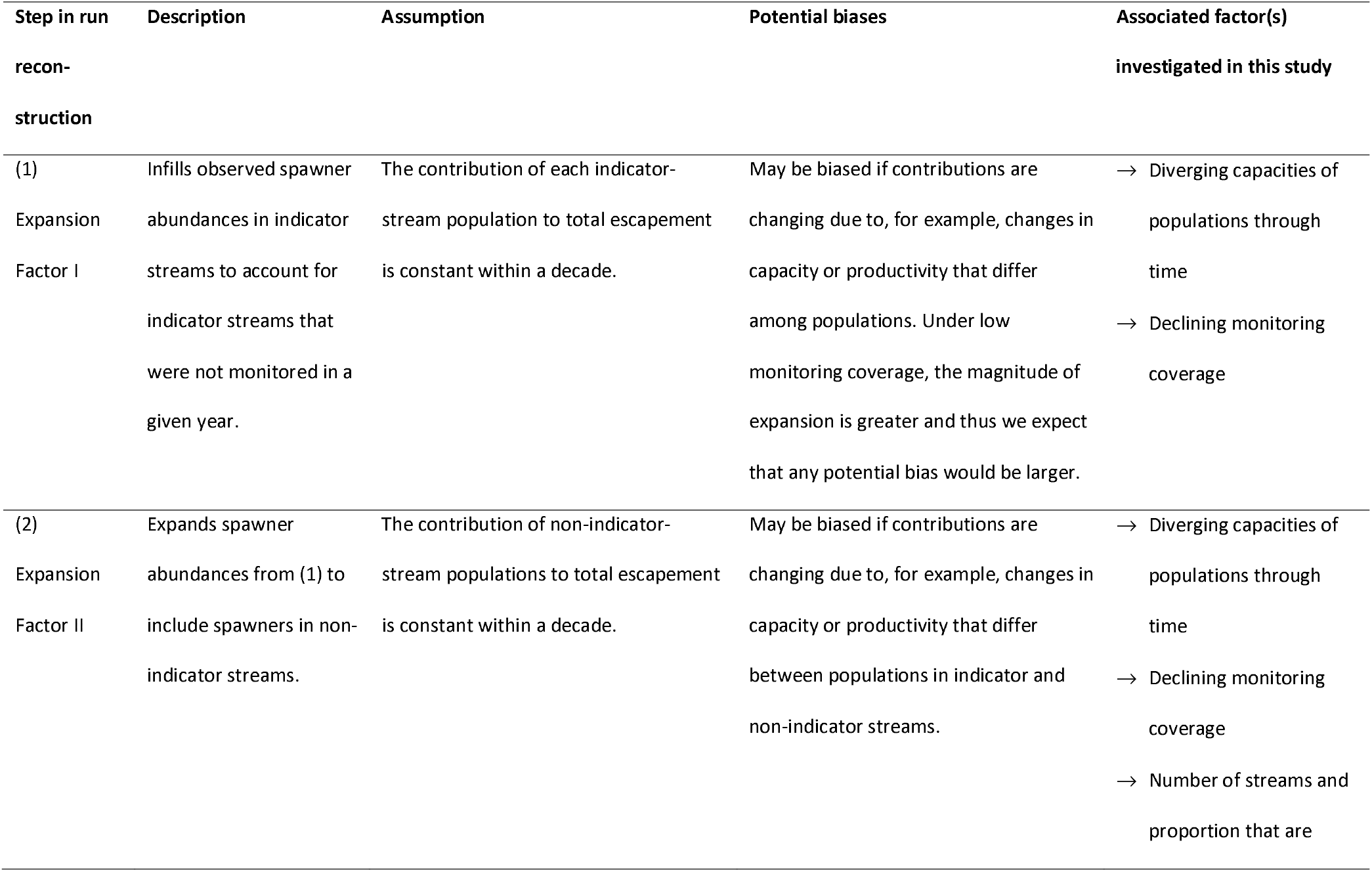
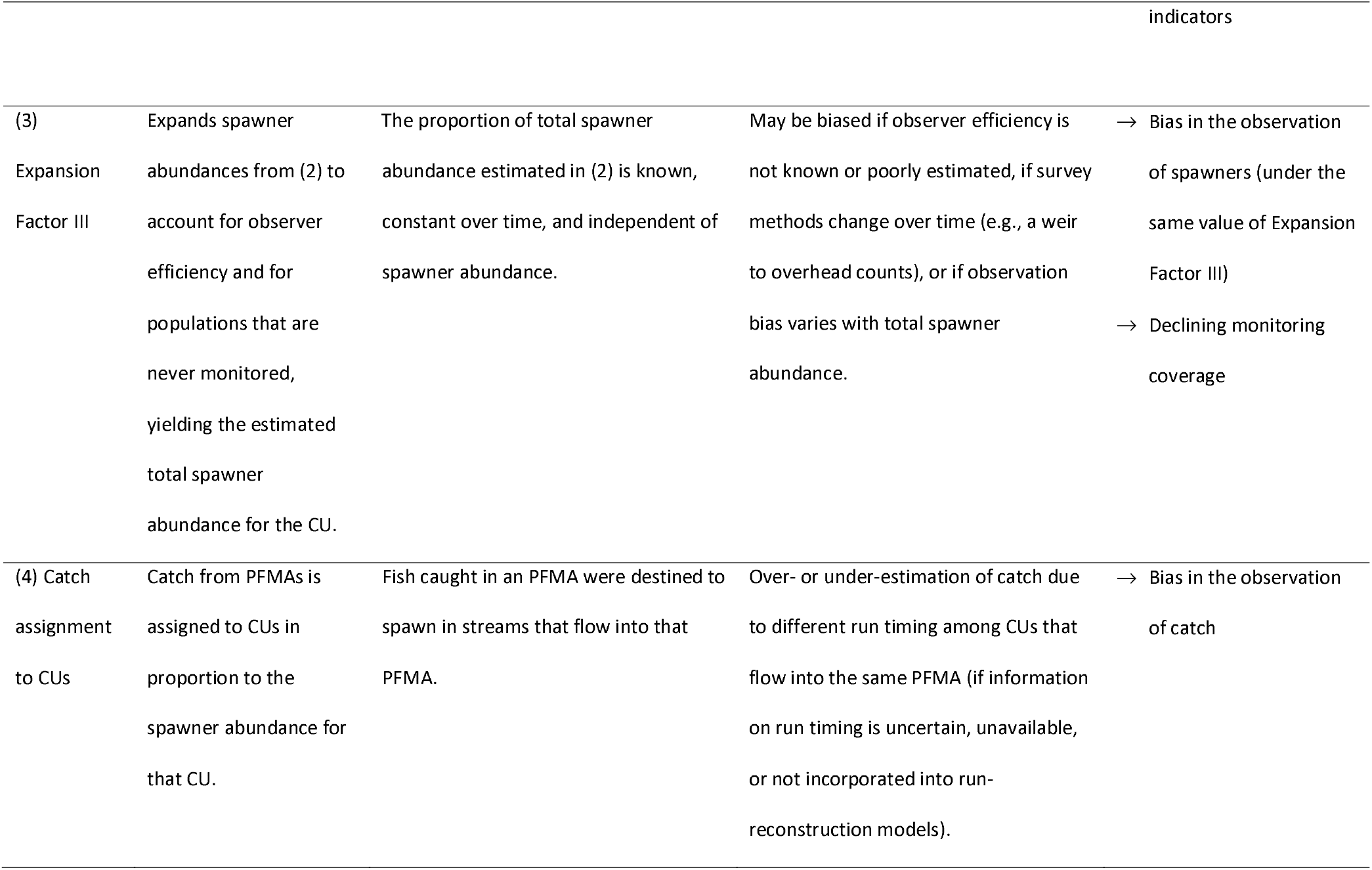
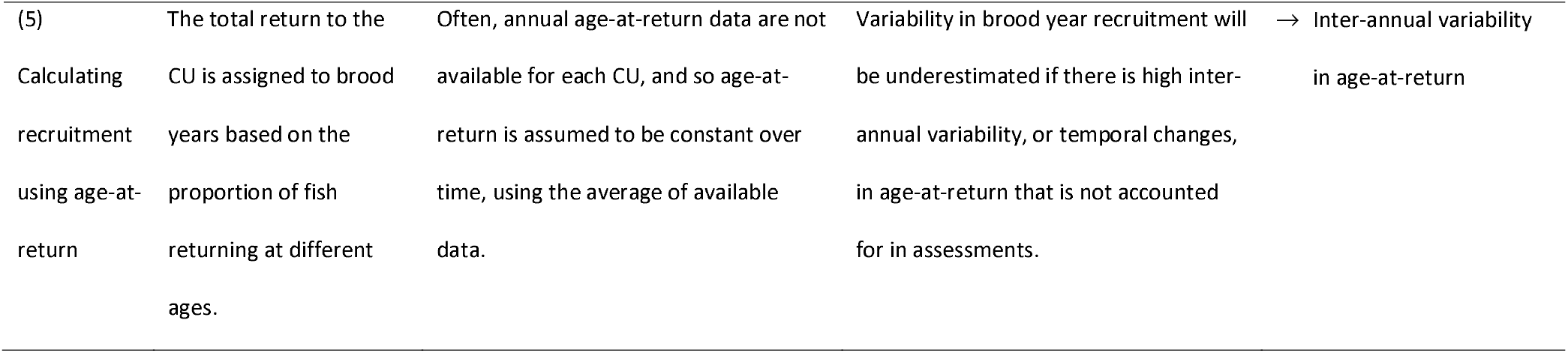
Summary of common steps in run reconstructions (Figure 2) and associated assumptions and potential biases that we investigated.

In this study, we quantified bias in biological status outcomes for Pacific salmon from data-driven assessments due to assumptions and uncertainty in run reconstructions (Table 1). We used a simulation approach that allowed us to explore how bias changed under different biological and management conditions. As a case study, we tailored our simulation model to represent a generic chum salmon CU from the central coast of BC because these CUs have recently been assessed using a data-driven approach (Connors et al. 2018) and have a relatively simple run-reconstruction model that does not include run timing. Furthermore, there are conservation concerns for both north and central coast chum salmon, which have not recovered despite significant reductions in harvest rates over the past two decades (DFO 2018b). Thus, central coast chum salmon offer a useful case study for an initial investigation of basic assumptions underlying biological status assessments. However, our simulation model is flexible enough to accommodate different species and life-history traits (e.g., age-at-return, density dependence) of Pacific salmon. We further explore a broad range of biological (e.g., trends in capacity, interannual variability in age-at-return) and management (e.g., declining monitoring coverage, bias in observed spawners and catch) scenarios to yield more general insight into the circumstances under which assumptions may bias status assessments.

## Methods

### Benchmark calculations and assumptions

Multiple metrics of biological status have been proposed under the WSP that cover four broad classes of indicators: current spawner abundance, trends in abundance over time, spatial distribution of spawners, and fishing mortality (Holt et al. 2009). We focused our analysis only on the current spawner abundance metric and considered two types of benchmarks (Figure 1,2) that have been frequently applied to determine biological status of Pacific salmon CUs, including in the Pacific Salmon Explorer (PSE; Connors et al. 2013, 2018, 2019). The first type of benchmark is associated with maximum sustainable yield, derived from a SR relationship (Figure 2b). An upper SR benchmark of 80% of the spawner abundance that is projected to maintain long-term maximum yield, or 80% *S*_MSY_, has been recommended by Holt et al. (2009, 2018) and will be applied to future assessments in the PSE (previous PSE assessments have applied *S*_MSY_; Connors et al. 2018, 2019). *S*_MSY_ can be calculated explicitly from the productivity and density-dependence parameters of the Ricker SR relationship (Scheuerell 2016). Multiple lower benchmarks have been suggested (Holt et al. 2009, 2018), and we used a lower benchmark of the spawner abundance that leads to *S*_MSY_ in one generation in the absence of fishing mortality, called *S*_GEN_ (Korman and English 2013; DFO 2015), as applied in the PSE. These benchmarks delineate the green, amber and red status zones: If the geometric mean spawner abundance over the most recent generation (*S*_AVG_) is above the upper benchmark of 80% *S*_MSY_, then the CU is assigned a green status, if *S*_AVG_ is between 80% *S*_MSY_ and *S*_GEN_ > then the CU is assigned an amber status, and if *S*_AVG_ is less than *S*_GEN_, then the CU is assigned a red status.

**Figure 2.**
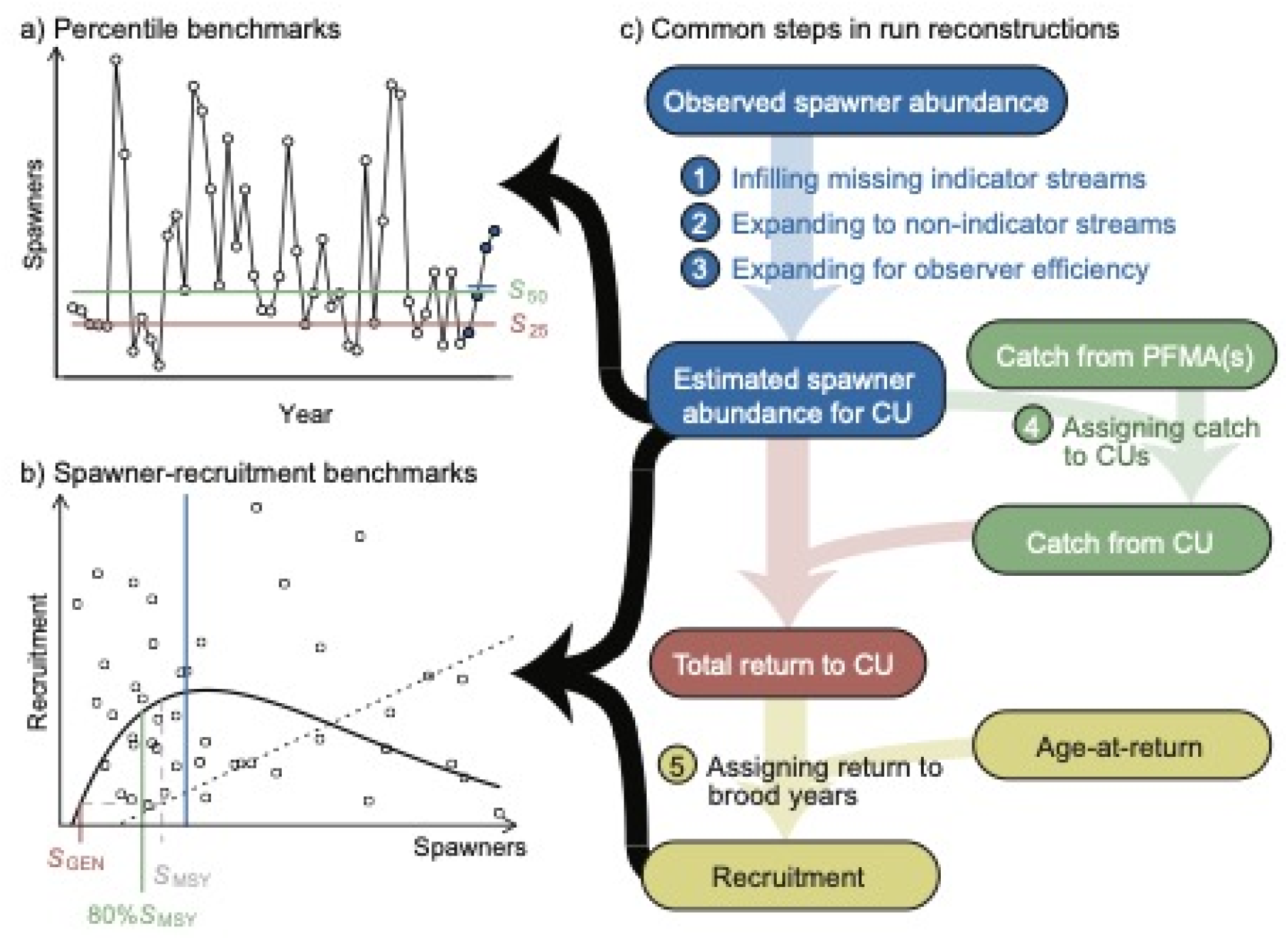
a) benchmarks are the 50^th^ (horizontal green line) and 25^th^ (horizontal red line) percentiles of historical spawner abundance (points). The current spawner abundance is calculated as the geometric mean spawner abundance over the most recent generation (4 years, blue points and line). b) -recruitment benchmarks are based on the shape of the Ricker relationship (solid line) fit to data on spawner abundance (x-axis) and corresponding recruitment (catch + escapement, y-axis). The upper and lower benchmarks are 80% S_MSY_ (green) and S_GEN_ (red), respectively. S_GEN_ is defined as the spawner abundance that leads to S_MSY_ (grey) in one generation in the absence of fishing mortality. Under both types of benchmarks, the current spawner abundance in the example shown is above the upper benchmark, and this CU would be assessed as ‘green’. c) The calculation of benchmarks and benchmarks requires run reconstruction to expand observed spawners abundances, assign catch to CUs, and calculate recruitment (Table 1).

The second type of benchmarks we considered were percentile benchmarks (Clark et al. 2014; Holt and Folkes 2015; Holt et al. 2018), also called historic spawners benchmarks (e.g., Connors et al. 2018). The upper and lower benchmarks are the 50^th^ and 25^th^ percentiles of historical spawner abundance, respectively (Figure 2a). Calculating these benchmarks requires fewer data, as the SR relationship need not be estimated (i.e. age-at-return and harvest data are not required). As such, the percentile benchmarks can be applied to data-limited CUs for which spawner data are patchy or age and harvest data are not available.

We focused on a basic run-reconstruction model and associated assumptions that are commonly made when assessing spawner abundance against the benchmarks above (Figure 2c; Table 1). At a minimum, in order to apply percentile benchmarks, a time series of total spawner abundance at the CU scale is required. Conservation Units are typically composed of multiple spawning populations that may or may not be monitored in any given year. Spawning populations in individual streams (henceforth “populations”) may exhibit unique dynamics as their productivity is (in part) limited by density-dependent processes in freshwater. A simple sum of the observed spawner abundance within a CU may be misleading if the same populations are not monitored consistently. On BC’s north and central coast, monitored populations have been designated as either “indicator streams” or “non-indicator streams”, with indicator streams being prioritized for monitoring and thus having more continuous and reliable spawner estimates (English 2016). In addition, there may be populations that have never been monitored and for which spawner abundance is unknown. To reconstruct spawner abundance to the CU, three “Expansion Factors” have been applied to account for (1) spawners returning to indicator streams that are not monitored in a given year, (2) spawners returning to non-indicator streams, and (3) observation efficiency and populations that are never monitored (Table 1).

The application of SR benchmarks also requires time series of the total number of salmon returning to the CU to reconstruct recruitment, including those caught in fisheries and those that make it to spawn but are not counted. The number of returning salmon in a CU that are caught in fisheries is estimated based on the catch statistics for Pacific Fisheries Management Areas (PFMAs) adjacent to the geographic location of the CU (Figure 3). It is assumed that salmon caught in a PFMA were destined to spawn in streams that empty into that PFMA, although there is the potential for bias in that fish may be caught while migrating through the PFMA or fish destined for streams in the focal PFMA may be caught in other PFMAs. Furthermore, in most cases, there is not a perfect spatial correspondence between PFMAs and CUs (Figure 3). Streams in multiple CUs may flow into a single PFMA, which is common for small CUs, such as with sockeye salmon which spawn in individual lakes, each of which is associated with a unique CU (Holtby and Ciruna 2007). In the simplest case, the catch from that PFMA may be assigned to CUs based on the relative spawner abundance to each CU. However, differences in run-timing among CUs may complicate the assignment of catch and necessitate more complex run-reconstruction models. A single CU may also be composed of populations that are caught in multiple PFMAs, particularly for species with large CUs such as pink and chum salmon whose geographic boundary often spans hundreds of kilometers and includes dozens of spawning populations (Figure 3). In these cases, an average harvest rate across PFMAs may be applied which may not reflect the variable harvest rates experienced by the different populations within these large CUs. The impact of observation bias in the catch assigned to each CU on status assessments is unknown and is a focal aspect of this study (see Sensitivity analyses, below).

**Figure 3.**
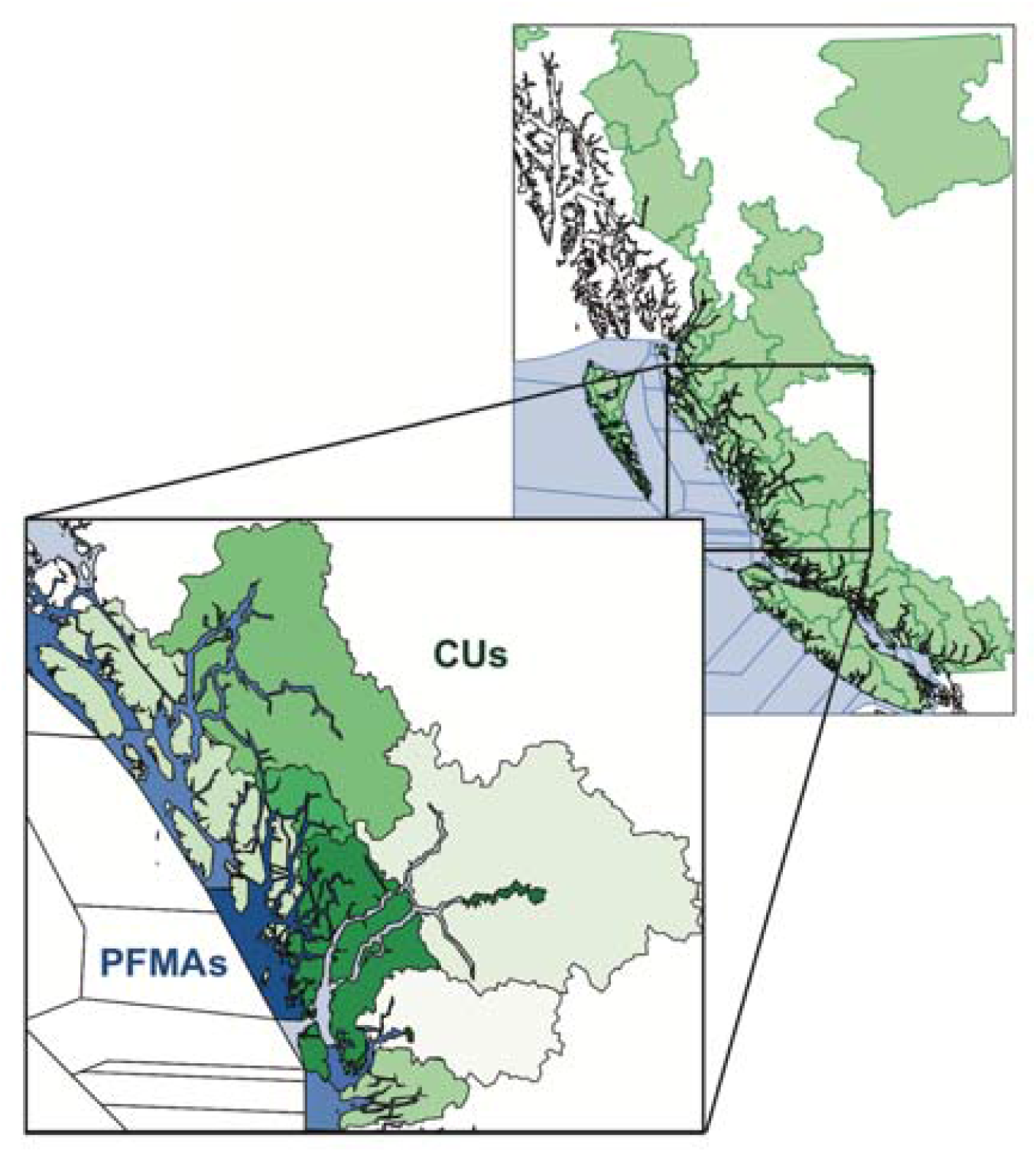
Our study focused on central coast chum salmon Conservation Units (CUs) (green; different CUs shaded differently in central coast inset), which are relatively large and do not correspond to the Pacific Fishery Management Areas (PFMAs; white or light blue shaded regions) for which catch is reported. One factor we investigated was how bias in assigning catch to CUs, resulting in under- or over-estimation of catch, affects estimates of biological status. Map produced using PBSmapping (Schnute et al. 2015) with data from Martin Huang (pers. comm.).

Finally, to calculate recruitment for a given cohort of spawners, assumptions about the age-at-return of spawners in any given year are required (except in the case of pink salmon, which have a fixed 2-year generation time). The total return in a given year is assigned to brood years based on the proportion of fish that return at a certain age, but these proportions are often not estimated every year. For chum salmon on the central coast, the distribution of age-at-return is assumed to be constant over time and is based on the average of available data (English et al. 2018). In this case, interannual variability in age-at-return may introduce uncertainty into the calculation of brood-year recruitment and bias resulting assessments of status (Zabel and Levin 2002).

### Simulation model

We developed and applied a stochastic simulation model of salmon population dynamics that allows control over various biological and management factors that may influence the accuracy of status assessments. Our approach built on previous studies that evaluated uncertainties in fisheries management (e.g., Holt and Peterman 2008) and other factors influencing the performance of metrics and benchmarks under the Wild Salmon Policy (e.g., Peacock and Holt 2012; Holt and Folkes 2015; Holt et al. 2016, 2018). The simulation model is composed of submodels for salmon population dynamics, observation of spawners, assessment, harvest, and performance (Figure 4). In this subsection, we describe the general equations for the simulation model, with details of parameterization in the following subsection. The code for the model and results is available at https://github.com/sjpeacock/run-reconst-sim_PSF. Version 1.0, used in this paper, is archived at doi: 10.5281/zenodo.3971276 and was implemented in R version 4.0.2.

**Figure 4.**
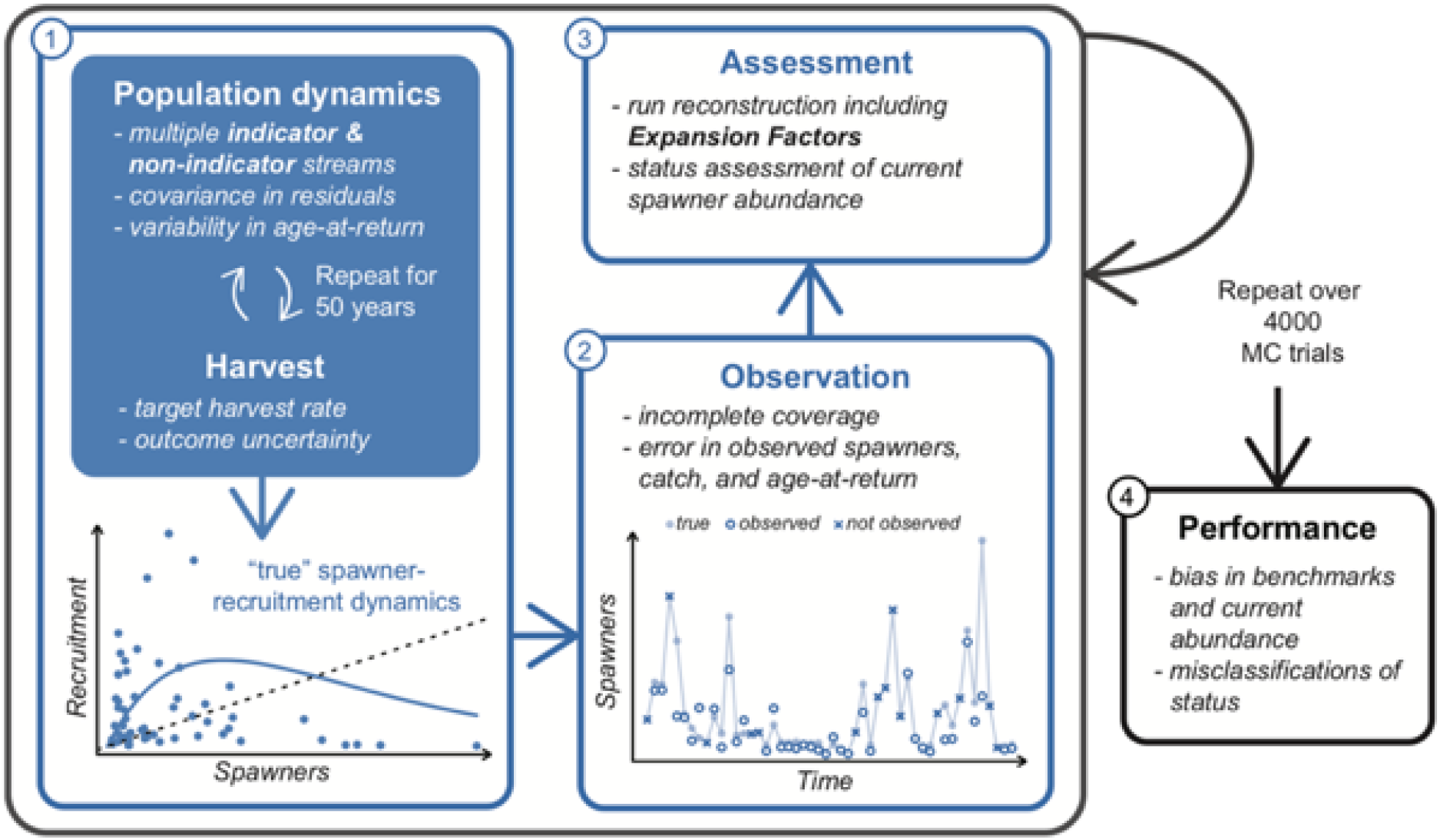
Schematic of the simulation model composed of submodels for population dynamics (including harvest), observation, assessment, and performance. Adapted from Holt et al. (2016).

#### Population dynamics

We simulated the dynamics of multiple spawning populations returning to indicator or non-indicator streams within a single hypothetical CU. Although some CUs consist of just a single spawning population (e.g., lake-type sockeye salmon), many CUs (especially pink and chum salmon) span thousands of square kilometers (Figure 3) and can include multiple spawning populations whose dynamics may differ due to local adaptation and finite rearing and spawning habitats.

We based our simulations on the life history of chum salmon, which generally return to spawn as 3-, 4-, or 5-year-olds. The number of salmon returning in return year *t* and population *j, R_t,j_*, was calculated as:

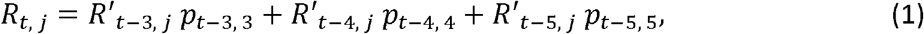

where *p_y,g_* is the proportion of recruits from brood year *y* returning at age *g*. Throughout this model description, we use *R* to denote returns, or catch plus escapement of fish returning in a year, and *R*’ to denote recruitment, or the total number of offspring from a brood year that survive to maturity.

We assumed that the annual proportion of recruits returning at a given age was the same among populations, but incorporated interannual variability in age-at-maturity by allowing the proportion of recruits that return at age *g* to vary among brood years, *y*.

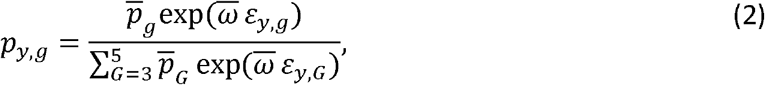

where 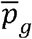 is the average proportion of individuals maturing as *g* year-olds, 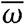 is a parameter that controls interannual variability in proportions of fish returning at each age (Figure S4) and *ε_y,g_* are standard normal deviates (Schnute and Richards 1995).

The number of salmon that escape the fishery and return to spawn was calculated as:

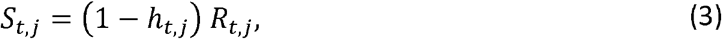

where *h_t,j_* is the realized harvest rate for population *j* in year *t*. We incorporated outcome uncertainty (i.e., deviations from the target harvest rates) by drawing the realized harvest rates for each year and population from a Beta distribution with mean equal to a target harvest rate, 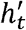, and variance 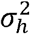 (Holt and Peterman 2008).

We considered two different scenarios for determining the target harvest rate (Figure 5). First, we considered a simple, abundance-based harvest control rule (HCR) where 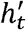 increased with the total return to the CU from a minimum of 0.05 (to account for bycatch and unavoidable mortality and also avoid problems associated with low target HRs when incorporating Beta-distributed outcome uncertainty) up to an asymptote, *h_MAX_* (Holt and Peterman 2008):

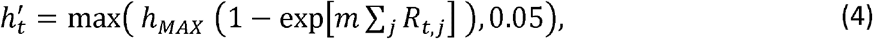

where *m* is the shape parameter of the HCR. The low harvest rates at low returns under this HCR prevented the CU from declining to red status in simulations, and so as to broaden our results to include CUs with true red status, we also considered a constant high target harvest rate of 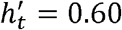 regardless of the total return (Figure 5). In the Supplementary Material, we present an intermediate scenario with a constant moderate target harvest of 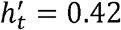.

**Figure 5.**
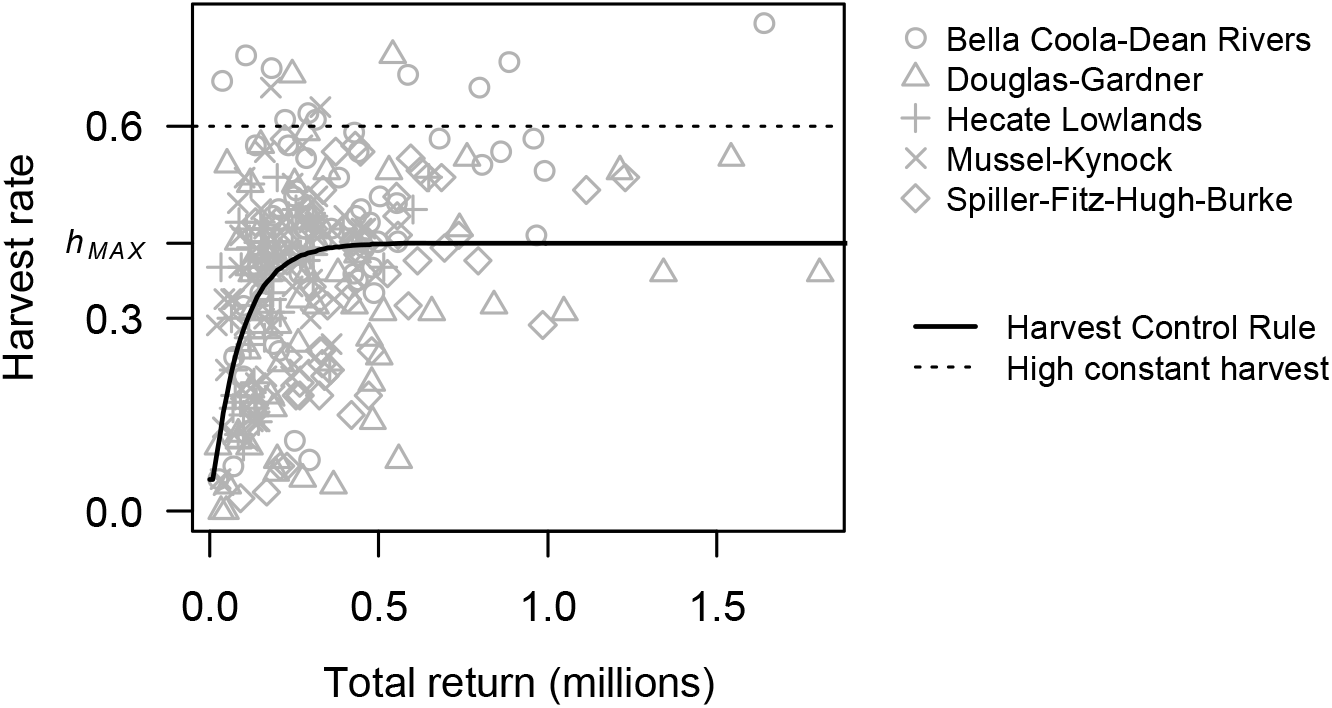
The two target harvest-rate cases we simulated were (1) a simple Harvest Control Rule (eq. (4); solid line) with parameters estimated from historical harvest rates and total return from five central coast chum CUs (grey points), (2) a constant high target harvest rate of 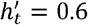 (dotted line).

Each population in our model was harvested in proportion to its abundance, such that the true total catch of fish that would have returned to streams within the CU was calculated as:

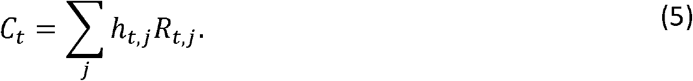

Although realized harvest rates differed among populations, we did not incorporate persistent biases in realized harvest rates among populations and thus assumed that all populations were equally vulnerable to the fishery. The extent to which this assumption is violated will depend on the size of the CU, the number of populations within it, the magnitude of variation in run-timing and body size among populations, and where fisheries are prosecuted. Any such biases among populations within a CU would likely be small because run-timing was a consideration when delineating CUs (Holtby and Ciruna 2007). However, biases among CUs may be significant and we investigate this by varying the observation bias in the total catch to the CU (see Sensitivity analyses, below).

Finally, we assumed the spawner-recruitment dynamics followed a Ricker relationship (Ricker 1954) yielding the number of recruits from brood year *y* and population *j*:

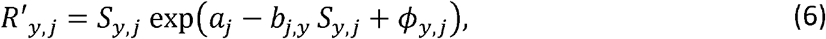

where *a_j_* is the log_*e*_ recruits per spawner at low spawner abundance (i.e., productivity), *b_j,y_* is the density-dependence parameter (which we allowed to change linearly through time; see below), and *ϕ_y,j_* are the recruitment deviates applied for year *y* and population *j* (eq. (7)).

We allowed productivity to differ among populations, where *a_j_* was drawn from a truncated normal distribution with mean *ā* and variance 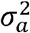, with a lower limit of *a_j_* = 0.4 because SR benchmarks are not calculable for very low productivity (Holt and Ogden 2013; Holt et al. 2018). For central coast chum salmon, we found only 1% of spawning populations (2/181) had *a_j_* < 0.4 (Online Supplement). A linear change in the density-dependence parameter was simulated for some populations as a decline in the capacity of the freshwater habitat (i.e., *S_MAX,j,y_* = 1/*b_j,y_*, or the spawner abundance that leads to maximum recruitment). This decline in capacities captured the potential consequences of cumulative stressors to freshwater habitat among watersheds on the central coast (Connors et al. 2018). For all populations, the initial capacity *S_MAX,j,1_* was drawn from a lognormal distribution whose mean and standard deviation differed for populations in indicator versus non-indicator streams, as indicator streams tend to be larger systems (English 2016; see Parameterization). The productivity and density-dependence parameters were drawn independently for each Monte Carlo (MC) iteration of the model.

We incorporated temporal autocorrelation in recruitment deviates:

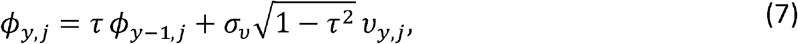

where *τ* is the temporal autocorrelation coefficient, *σ_v_* is the standard deviation in recruitment deviates without temporal autocorrelation (Ricker 1975, Holt and Bradford 2011), and *ν_y,j_* is a multivariate normal random variable with mean zero and variance-covariance matrix:

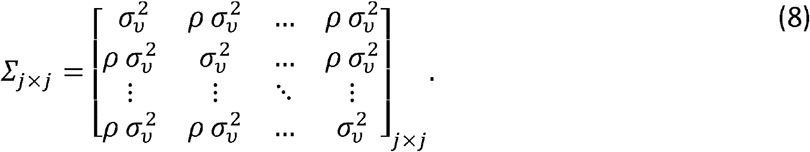

Here, *ρ* is the correlation in recruitment deviates among populations.

We simulated the ‘true’ population dynamics over 50 years, after an initialization period of seven years to seed eq. (6) given the variable age-at-return of chum salmon. For each year in this initialization, we assumed that the number of spawners was equal to 20% of *S_MAX,j,1_*. For the first year of the initialization, we set *ϕ_y−1,j_* from eq. (7) to zero.

#### Observation submodel

In the observation submodel, we incorporated both incomplete monitoring coverage of streams and imperfect observation of spawners in streams that were monitored. In any given year, population *j* was observed with probability *ψ_t,j_*. We included a linear decline in monitoring coverage (i.e., the probability of a population being observed) over time based on observations of declining monitoring coverage on the north and central coast of BC (Price et al. 2008, 2017; English 2016). We calculated the annual probability of being monitored separately for indicator and non-indicator streams based on observations that monitoring coverage of non-indicator streams is generally lower and has declined more severely than coverage of indicator streams (English 2016). See Parameterization for further details.

Spawner abundances were ‘observed’ with log-normal error:

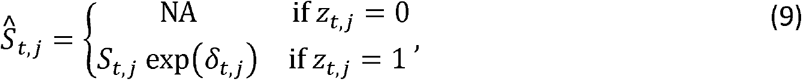

where 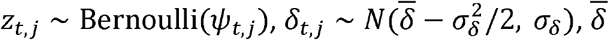 is the mean observation error, and *σ_δ_* is the standard deviation in observation error of spawner abundances. Thus, this combines both the probability of a population being monitored and the distribution of observation errors around true spawner abundances if monitored. The mean observation error is corrected by 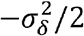 so that the arithmetic mean observation bias is 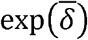. We included a negative bias in the observation of spawners 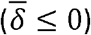 such that the observed spawner abundance is on average lower than the true spawner abundance. In general, it is challenging to enumerate spawners in all reaches of a stream and in all streams within a CU. The reported spawner abundance is considered an underestimate of the total spawners in a CU, which motivates the application of Expansion Factor III for observer (in)efficiency when performing run reconstructions (Table 1). The calculation of Expansion Factor I required that we impose the constraint that at least one indicator stream was monitored each year, so if *z_t,j_* = 0 for all indicator streams in a year, we randomly selected one indicator stream to be monitored.

The catch to the entire CU in return year *t* was observed with log-normal error:

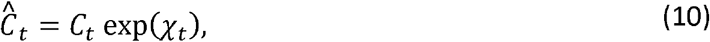

where 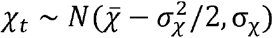, *σ_χ_* is the standard deviation in catch error, and 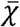 is a bias in catch (corrected by 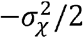 as for observed spawners above). We assumed a default value of 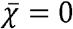, but in sensitivity analyses we varied this parameter to simulate scenarios in which fish are caught from other CUs or fish from the focal CU were caught in other PFMAs.

Previous models (e.g., Holt et al. 2018) have included error in the “estimated age-at-return” separately for each return year. For central coast chum salmon, annual age-at-return data are rarely sampled comprehensively so the same average is generally applied across all years (Peacock et al. 2014; English et al. 2018). Therefore, for each year in a MC trial we applied the same age-at-return, which was drawn independently for each MC trial using eq. (2) with observation error, 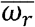. (Table A1).

#### Assessment submodel

As described above, the minimum requirement to calculate benchmarks and assess population status using percentile benchmarks is a time-series of spawner abundance to the CU. For SR benchmarks, harvest rates and age-at-return must also be estimated in order to calculate recruitment. The basic procedure of these run reconstructions is outlined in Table 1, and begins by expanding observations of spawners to indicator streams to the total spawner abundance to the CU by applying three Expansion Factors. The equations and criteria governing these Expansion Factors are detailed in the Online Supplement and in English et al. (2012, 2016, 2018). Briefly, Expansion Factor I 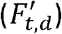 imputes missing spawner abundances for populations in unmonitored indicator streams and is calculated for each year, *t*, within a decade, *d*, of the spawner time series. It relies on the decadal contribution of each indicator-stream population to the total escapement to all indicator streams (English et al. 2016). Expansion Factor II 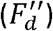 expands observations of spawners from indicator streams to include populations in non-indicator streams that are less frequently monitored, and is the same for each year within a decade, *d*. Expansion Factor II is calculated as the average proportion of total monitored spawners (in indicator and non-indicator streams) that are in non-indicator streams for that decade. For decades with insufficient information to calculate either of these Expansion Factors, for example due to declining monitoring coverage, a reference decade may be used. Expansion Factor III (*F*”’) is determined by the regional DFO staff familiar with the escapement monitoring techniques used in each statistical area and is assumed to be constant through time (English et al. 2018). In our model, we assumed that all populations were at least partially monitored, and that Expansion Factor III accounted for observation (in)efficiency, but in reality, Expansion Factor III may also account for populations in unmonitored streams.

The observed number of salmon returning in year *t* is the sum of observed catch and expanded escapement to the CU:

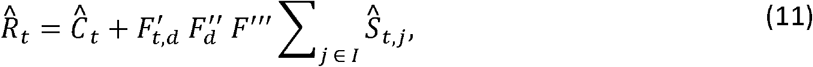

where the summation includes observed spawner abundance to the *I* indicator streams only, with the non-indicator streams being accounted for through Expansion Factor II, 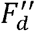.

We do not explicitly account for en route or pre-spawning mortality of fish that escaped the fishery, and assume that pre-spawning mortality is relatively small and accounted for in the productivity of the population through the Ricker SR dynamics. Observed recruitment for brood year *y* is calculated as the sum of age 3, 4, and 5 fish returning in years *y* + 3, *y* + 4, and *y* + 5, respectively:

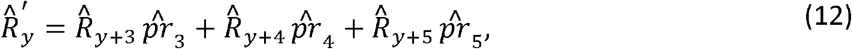

yielding the ‘reconstructed’ spawner-recruit pairs for brood year *y*.

To calculate estimated SR benchmarks, we fit a linearized Ricker model to the observed data at the aggregate CU-level. The estimated productivity and density-dependence parameters were then used to calculate upper and lower SR benchmarks (80% of *S_MSY_* and *S_GEN_*) for the CU.

#### Performance submodel

For each MC simulation, we estimated status using both SR and percentile benchmarks calculated from the observed spawner-recruit pairs for the CU, including observation biases and incomplete monitoring coverage. Estimated status under both types of benchmarks was compared to the true status, which was calculated by comparing the current true spawner abundance (without observation error) against the upper and lower SR benchmarks (80% of *S*_MSY_ and *S*_GEN_) from the underlying SR parameters. This meant that we evaluated estimates of percentile (and SR) benchmarks against the true underlying benchmarks derived from “true” SR parameters. Because we simulated the true dynamics at the scale of spawning populations and there was no “true” CU-level value for the Ricker parameters, calculating the true *S*_MSY_ (and thus the SR benchmarks) at the CU-level was not straightforward. We chose to calculate the true SR benchmarks for each population from the underlying Ricker parameters for that simulation and then summed the benchmarks across all populations to yield the “true” CU-level benchmarks. Although this CU-level benchmark will underestimate the level required to maintain all component populations above their individual benchmarks in any given year, the objective of the WSP is to maintain the overall CU and populations within CUs are assumed be recolonizable within reasonable time frames (Fisheries and Oceans Canada 2005). When declines in capacity were included in the simulation, we calculated true status from the initial capacity parameters before the decline. Although this may have increased the chance that estimated status was lower than true status (increasing the rate of pessimistic misclassifications), we took this approach to avoid a shifting baseline in benchmarks.

Performance was evaluated in two ways that capture the difference between estimated and true status: (1) the proportion of MC simulations for which status was correctly assessed as green, amber, or red, and the proportion of simulations for which status was either underestimated (pessimistic) or overestimated (optimistic), and (2) the percent relative bias of observed average spawner abundance (*S*_AVG_) and of the four benchmarks (*S*_GEN_, 80% *S*_MSY_, *S*_25_, *S*_50_) compared to their true values for each MC simulation.

For each parameterization investigated (see below) we ran 4000 MC trials, which was sufficient to ensure the mean percent error in performance measures was < 3% (Figure S3).

### Parameterization

Some of the parameters in our simulation model were unknown or unknowable, in which case we followed assumptions made for southern BC chum salmon by Holt et al. (2018). Other parameters were available specifically for central coast chum salmon or could be estimated from available data; details of parameter estimation are given in the Online Supplement. As mentioned above, in order to understand the assessment biases under different true statuses we considered two cases: (1) high productivity and a conservative harvest control rule (HCR), which we refer to as the “base case” because it is most representative of central coast chum salmon, and (2) low productivity and a constant high target harvest rate, which represented a CU at high risk of extirpation. In the Online Supplement, we also present results from a third case intermediate between these two with low productivity and a constant moderate harvest rate. Unless otherwise indicated in the sensitivity analysis (below), parameters defaulted to the values described here and listed in the Appendix, Table A1.

The mean proportion of adults maturing at ages 3, 4, and 5 in eq. (2) was held constant for all simulations at 0.23, 0.64, and 0.13, respectively, based on the average age-at-return applied in run reconstructions of central coast chum (Challenger et al. 2018; English et al. 2018). For the base case, we estimated the parameters in the HCR (Figure 5) from harvest rates and total return sizes for five central coast chum CUs (English et al. 2018; Salmon Watersheds Program - Pacific Salmon Foundation 2019). The high harvest case is consistent with the upper bound in harvest rates chosen in other studies (Holt et al. 2018) and represents the upper threshold in harvest rates for central coast chum (Figure 5).

To estimate the SR parameters for spawning populations (eq. (6)), we fit a linearized Ricker model to population-level spawner-recruit pairs from nine central coast chum CUs with individual productivity and density-dependence parameters for each population (Figure S5). Note that this approach may lead to biased SR parameters as we did not account for temporal autocorrelation and errors in variables when analysing the data, but they nonetheless provide a useful starting point for our simulation study. From the model fits, we calculated (1) mean and variance in productivity among populations, (2) mean and variance in the initial capacity (i.e., 1/*S*_MAX_) for indicator and non-indicator streams, (3) the residual variance within populations, (4) the average correlation in residuals between populations, and (5) the temporal autocorrelation in residuals within populations. The mean population-level productivity was *ā* = 1.40 (Figure S5), which we applied in our base case. For the low-productivity case, we chose *ā* = 0.56, which was the 2.5^th^ percentile of population-level productivity estimates (Figure S5). The residual variance within populations was used as an estimate of the variance in recruitment deviations 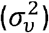 in simulations, but was likely an overestimate as we did not account for observation error separately when fitting the Ricker model. Thus, we explore the results under a range of values for *σ_v_* in the Supplemental Results (Online Supplement). The correlation in residuals among populations was estimated at *ρ* = 0.46, but we also investigated lower (*ρ* = 0) and higher (*ρ* = 0.9) levels of synchrony in the Online Supplement.

As the default case, we considered a decline in the capacity of streams (i.e., one over the density-dependence parameter of the Ricker model) that reflected observed declines in freshwater habitat (Office of the Auditor General of Canada 2004). Within the central coast chum CUs, 29% of watersheds are considered to be at moderate risk, and 21% at high risk, of cumulative habitat pressures over the last 60 years from stressors such as logging, water licenses for withdrawal of water from streams, and stream crossings (Connors et al. 2018). We hypothesized that declines in capacity that differ among spawning populations may affect the accuracy of Expansion Factors I and II, particularly in combination with declining monitoring coverage (Price et al. 2017). We incorporated a linear decline in capacity (i.e., *S*_MAX_ = 1/*b*) over the 50-year time series of between 25% and 50%, representing a moderate decline, for 29% of populations (the percentage of central coast chum watersheds deemed to be at moderate risk of cumulative habitat pressures by Connors et al. 2018) and a linear decline between 50% and 75%, representing severe decline, for 21% of populations (the percentage deemed to be at high risk). The exact percent decline for each population was randomly drawn from a uniform distribution within the above range for each MC simulation. The remaining 50% of populations had stable capacity over the 50-year simulation. In a sensitivity analysis, we investigated four additional scenarios for declining capacity (Table 2).

**Table 2.**
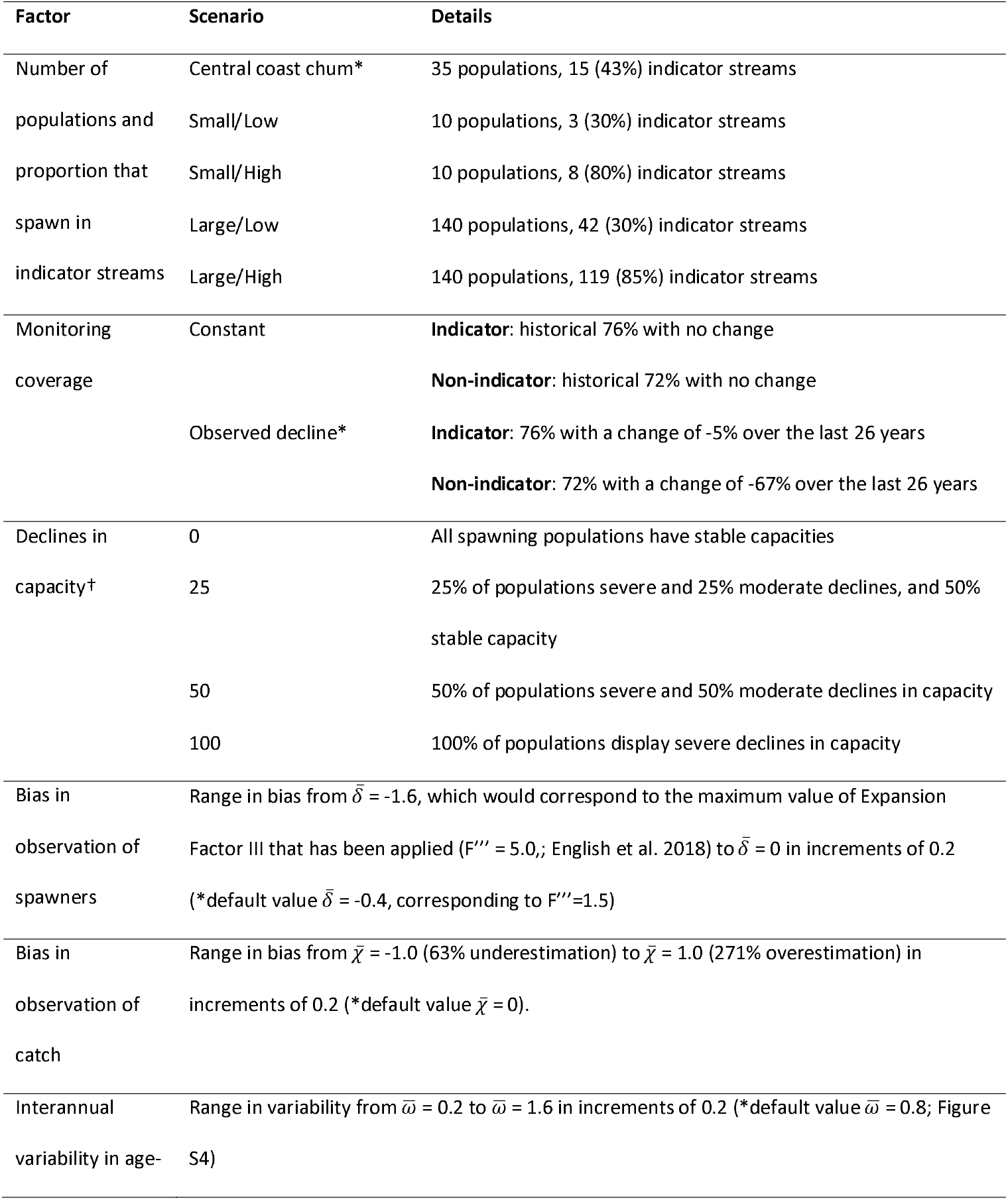
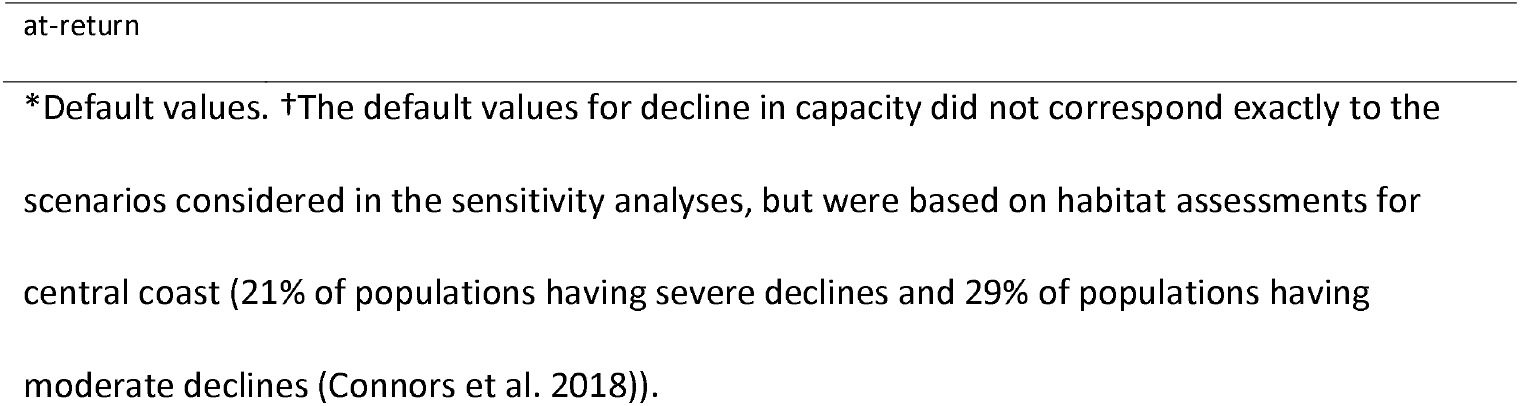
Summary of factors that we investigated in sensitivity analyses to determine their impact on bias in status assessments.

In the observation submodel, we chose the bias in the observation of spawners to match Expansion Factor III, which corrects for observer efficiency, with a range of values explored in a sensitivity analysis (below). Because Expansion Factor III is simply a constant multiplier applied to spawner abundance, it does not affect estimated status by the percentile benchmarks. However, Expansion Factor III affects estimated harvest rates and recruitment, and therefore bias in this Expansion Factor was expected to lead to bias in estimated status under the SR benchmarks. The value of Expansion Factor III applied in past status assessments has been constant at F’” = 1.5 for all central coast chum CUs (English et al. 2016), and so we applied a default value of 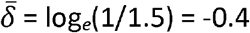. The majority of central coast chum streams are surveyed visually by ground (fish counts or other sampling) with some aerial (fish counts) or boat (fish counts or other sampling) surveys (English 2016), similar to southern BC chum streams (Holt et al. 2018). We assumed *σ_δ_* = 0.5 following Holt et al. (2018), which is the maximum estimated uncertainty for visually surveyed spawners (Cousens et al. 1982; Szerlong and Rundio 2008).

We incorporated a linear decline over the last 27 years of simulations in the proportion of indicator and non-indicator streams monitored each year from 0.76 and 0.72, respectively, to 0.72 and 0.05 based on English (2016). These declines are representative of overall declines in monitoring across species, but we also consider the trends specific to chum salmon in the Online Supplement (Figure S6).

We assumed no bias in the observation of catch 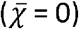 by default, but consider a range of bias in sensitivity analyses (below). The standard deviation in the observation error of catch accounts for differences between observed (i.e., reported) and actual catch due to uncertainties with mixed-stock fisheries and in reporting and estimation of recreational fisheries and subsistence use. We set this to *σ_χ_* = 0.2 (Holt et al. 2018), which is less than the observation error in spawners.

#### Sensitivity analyses

We examined the sensitivity of status assessments over a range of values for several different model parameters that we considered most likely to affect status assessments due to their influence on the assumptions in run reconstructions (Table 1; Table 2). We investigated each of the questions below under both the base case and low-productivity high-harvest case described above, with other parameters at their default values (Table A1) unless otherwise noted. The specific questions that we addressed were:

1. How does the number of spawning populations and the proportion designated as indicator streams affect status assessments? The lower the proportion of streams that are indicators, the greater the magnitude of Expansion Factor II.
2. How does a decline in monitoring coverage affect status assessments? The fewer indicator streams that are monitored, the greater the magnitude of Expansion Factor I and the potential uncertainty in expanded spawner abundance. Here, we consider two scenarios (Table 2; Figure S6): constant monitoring coverage at historical proportions among all streams and an observed decline in coverage starting in the mid-1980s as has been observed on the north and central coast (English 2016; Figure S6). In the Online Supplement we consider two additional scenarios: observed declines in monitoring specific to chum salmon streams and a sharp, recent decline in monitoring of indicator streams.
3. How do declines in capacity affect status assessments? The application of Expansion Factors I and II assumes that the relative contributions of populations to aggregate abundance in the CU does not change over time, but declines in capacity that differ among populations may violate this assumption.
4. How does spawner observation bias affect status assessments, given that the value of Expansion Factor III is fixed over time and often the same among CUs (English et al. 2018)?
5. How does catch observation bias (e.g., over- or under-estimating catch of salmon) affect status assessments? This represents scenarios where there are errors in estimates of CU proportions in the aggregate catch in a mixed-stock fishery, or violation in the assumption of homogenous spatial and temporal distribution of CUs when CU proportions are not monitored in such fisheries.
6. How does interannual variability in age-at-return affect status assessments?

We investigated the impact of declines in monitoring coverage (question #2 above) in combination with declines in capacity of spawning populations (question #3) in a bivariate sensitivity analysis.

## Results

The different productivity and harvest rate combinations we considered led to different true CU statuses. Under high productivity and an abundance-based harvest control rule (HCR) – the base case corresponding to central coast chum salmon – 86.4% of simulations resulted in true green status (Figure 6a,b). Conversely, under low productivity and high harvest rates, 67.0% of simulations resulted in true red status (Figure 6c,d).

**Figure 6.**
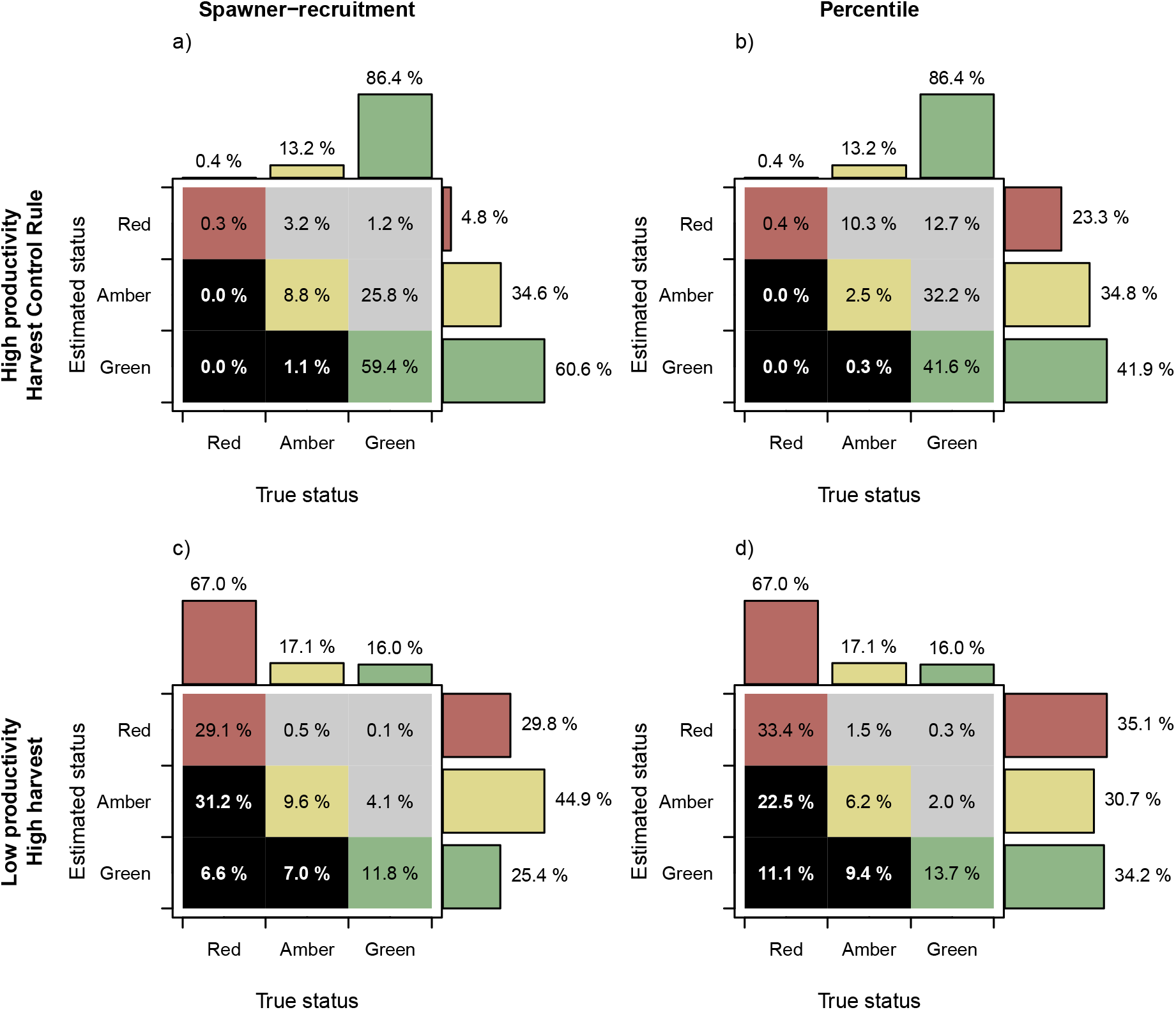
Estimated status according to the spawner-recruitment (SR) benchmarks (left) and the percentile benchmarks (right), over the true status for each of two cases (high productivity and an HCR (a-b) and low productivity and high harvest rates (c-d)). Grey cells indicate pessimistic misclassifications, which may lead to overly conservative management actions, and black cells indicate optimistic misclassifications, which may lead to overly risky management actions. Coloured cells indicate correct classifications for red, amber, and green zones, respectively.

Under the base case when true status was mostly green, misclassifications resulted in estimated status lower than the true status meaning assessments were biologically pessimistic (henceforth referred to as “pessimistic misclassifications”). This was particularly true of the percentile benchmarks, for which 55.2% of simulations resulted in a pessimistic misclassification with 12.7% of simulations having misclassified green status as red. Pessimistic misclassifications were due to positive bias in benchmarks and not bias in the current spawner abundance (Figure 7), resulting in status being underestimated. For productive populations (as in the base case), most observed spawner abundances tended to be far above lower benchmarks and closer to equilibrium values. As a result, lower and upper benchmarks of 25^th^ and 50^th^ percentile of historical spawner abundance tended to overestimate the “true” SR-based benchmarks.

**Figure 7.**
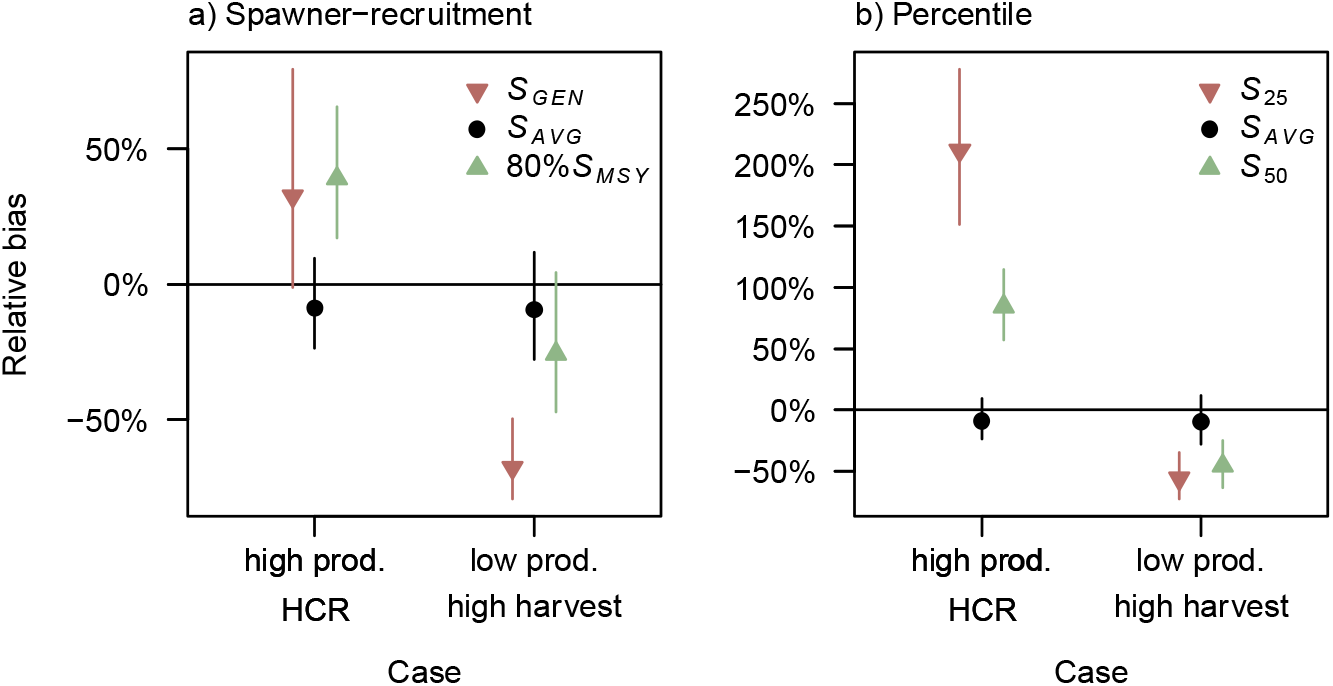
Relative bias in spawner-recruitment (SR) benchmarks (a), percentile benchmarks (b), and current spawner abundance (S_AVG_; black) (median ± interquartile range among 4000 MC simulations) for high productivity and a harvest control rule (HCR) and the low-productivity high-harvest case.

When true status was mainly red, under low productivity and high harvest rates, biologically optimistic misclassifications (henceforth “optimistic misclassifications”) were more common, which may be riskier from a conservation and management standpoint. For example, 44.8% and 43.0% of simulations had an estimated status higher than true status under the spawner-recruitment (SR) and percentile benchmarks, respectively (Figure 6c,d). These more frequent optimistic misclassifications were due to a negative bias in benchmarks, in particular the lower SR benchmark of *S*_GEN_ (Figure 7a), likely due to a poor ability to estimate *S*_MSY_ under low productivity when spawner abundances tend to cluster near the origin.

Under both types of benchmarks, bias did not decrease when monitoring coverage was held constant at 100% (Figure S9), suggesting that the application of Expansion Factors I and II were not contributing factors.

### Sensitivity analyses

The number of spawning populations within the CU and the proportion of those populations spawning in indicator streams had little impact on status assessments (Figure S10). Under the base case, the relative bias in estimates of *S*_MSY_ and *S*_GEN_ were lower in larger CUs, with half as many pessimistic misclassifications for larger CUs under the SR benchmarks (43% for 10 populations versus 22% for 140 populations with 30% indicator streams; Figure S10a). This trend was not, however, observed in the low-productivity high-harvest case when true status was predominantly red (Figure S12).

The monitoring-coverage scenarios that we considered, representative of observed declines in monitoring on the north and central coast, had no effect on status outcomes or the relative bias in benchmarks. This was true in the base case (Figure 8, Figure S13) and under low productivity (Figures S14 – S15). Even under severe declines in capacity of 50% to 75% for all spawning populations, our results suggest that the observed declines in monitoring coverage on the north and central coast are unlikely to bias status assessments. This result held regardless of whether the recruitment deviates among component populations within the CU were not correlated (*ρ* = 0; Figure S16) or highly correlated (*ρ* = 0.9; Figure S17).

**Figure 8.**
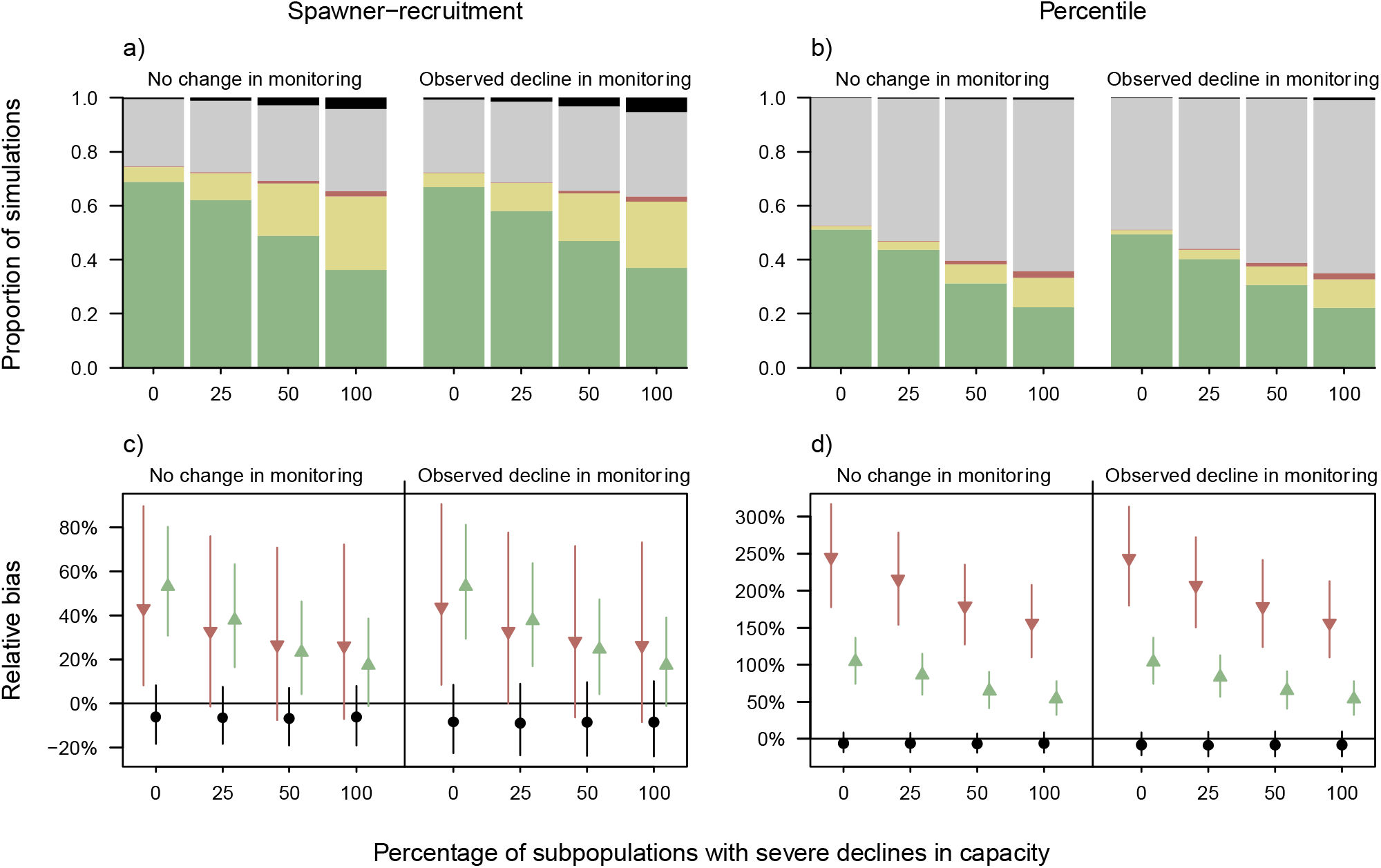
The effect of monitoring coverage (no change and decline; Table 2) and the percentage of spawning populations with severe declines in capacity (x-axis) on performance measures under the base case of high productivity and HCR. (a-b) The proportion of simulations with correct green, amber, or red status or pessimistic misclassifications (grey) and optimistic misclassifications (black) under the SR benchmarks (a) and percentile benchmarks (b). (c-d) The percent relative bias (median ± interquartile range among 4000 MC simulations) in the current spawner abundance (S_AVG_; black) and lower and upper benchmarks (red and green, respectively) under the SR benchmarks (c) and the percentile benchmarks (d). See Online Supplement for results under the low-productivity high-harvest case.

Under the base case, declines in capacity of the CU were associated with poorer estimated status and an increase in misclassification rates (Figure 8a-b). Pessimistic misclassifications increased because CUs more frequently had a true status of amber but were misclassified as red. Optimistic misclassifications increased, particularly under the SR benchmarks (Figure 8a), because the relative bias in the current spawner abundance (*S*_AVG_) remained unchanged, but the bias in benchmarks decreased (Figure 8c-d). In the low-productivity high-harvest case, the results were similar but with biologically optimistic misclassifications dominating as status was predominantly amber or red (Figure S15).

As the bias in the observation of spawners approached zero 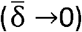, misclassifications under the SR benchmarks declined in all cases, even as the observation bias became less than the Expansion Factor III applied to correct for it (Figure 9a, Figures S18-S19). The relative bias in the current spawner abundance was minimized when the observation bias matched the assumed value of Expansion Factor III (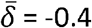 corresponding to F”’ = 1.5; Figure 9c-d). When observed spawner abundance was biased low 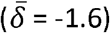, the relative bias in the upper benchmark of 80% *S*_MSY_ was higher than the relative bias in current spawner abundance (*S*_AVG_) or the lower benchmark (*S*_GEN_), and thus CUs with a true green status were more likely to be misclassified as amber. Status outcomes under the percentile benchmarks were unaffected by changes in observation bias of spawners (Figure 9b); the bias in both benchmarks and current abundance showed similar changes as observation bias declined (Figure 9d) such that the resulting status outcome was unchanged. In the Online Supplement, we also investigated a change in observation bias halfway through the simulation (Figure S20), but a change from the base value of 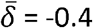 to 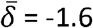, −0.7, and 0 did not have any impact on status outcomes or biases in benchmarks (Figure S21).

**Figure 9.**
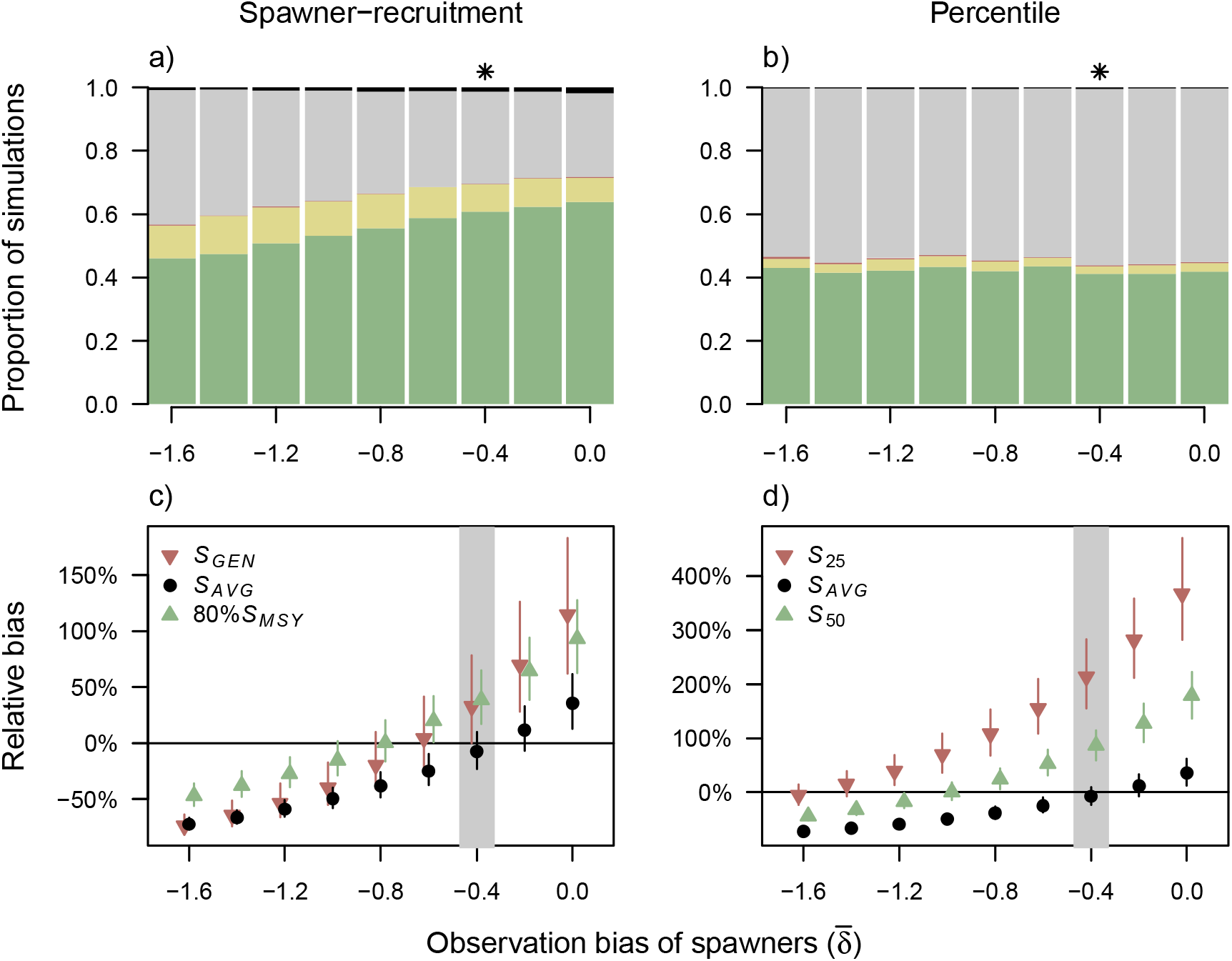
The effect of observation bias in the number of spawners (x-axis) on performance measures under the base case. (a-b) The proportion of simulations with correct green, amber, or red status or pessimistic misclassifications (grey) and optimistic misclassifications (black) under the SR benchmarks (a) and percentile benchmarks (b). (c-d) The percent relative bias (median ± interquartile range among 4000 MC simulations) in the current spawner abundance (S_AVG_; black circle) and lower and upper benchmarks (red and green, respectively) under the SR benchmarks (c) and the percentile benchmarks (d). The asterisk in (a-b) and grey zone in (c-d) indicate the default parameter value of 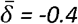, and the bias that matches the Expansion Factor III of F’” = 1.5 applied in all simulations. See Online Supplement for results under the low-productivity high-harvest case (results were similar).

Underestimation of catch (i.e., negative values of 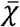) resulted in fewer misclassifications than overestimating catch (Figure 10, Figures S22 – S23). As the catch bias increased from 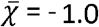 (63% underestimation) to 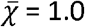 (271% overestimation), the relative bias in the lower SR benchmark of *S*_GEN_ declined while the relative bias in the upper benchmark of 80% *S*_MSY_ increased (Figure 10b). This is due to the errors in variables that occur when catch is underestimated: productivity and recruitment tend to be underestimated, thus leading to lower estimates of *S*_MSY_ and higher estimates of *S*_GEN_ (Holt and Folkes 2015). Under the base case, the true status was green in the majority of simulations and so the increasing bias in the upper benchmark dominated the overall status assessments and led to the increase in pessimistic misclassifications with increasing 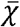. In the low-productivity high-harvest case, true status was mostly red and so the increasingly negative bias in *S*_GEN_ resulted in more optimistic misclassifications as the observation bias in catch increased (Figure S23). In all cases, overestimating catch by ~50% (i.e., 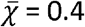) led to a 5 – 8% increase in misclassification rate (Figure S24). Although these changes in misclassification rates may seem small, there is potential for large catch errors in run reconstructions, especially when multiple CUs overlap with a single PFMA. Catch does not factor into the calculation of percentile benchmarks, so status under the percentile benchmarks was unaffected by changing catch bias.

Finally, increasing interannual variability in age-at-maturity resulted in more frequent status misclassifications, but the effect was relatively small. Under the base case, increasing 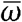 from 0.2 to the default value of 0.8 resulted in an increase in misclassifications from 27.3% to 30.2%, but very little change in the bias in benchmarks (Figure S25). Further increasing the interannual variability to 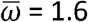 led to 34.0% of simulations being misclassified, but this level of interannual variability is high (see Figure S4 for example) compared to data on age-at-return for central coast chum salmon (Challenger et al. 2018; English et al. 2018). The increase in misclassifications was smaller under the low-productivity high-harvest case (Figure S25).

**Figure 10.**
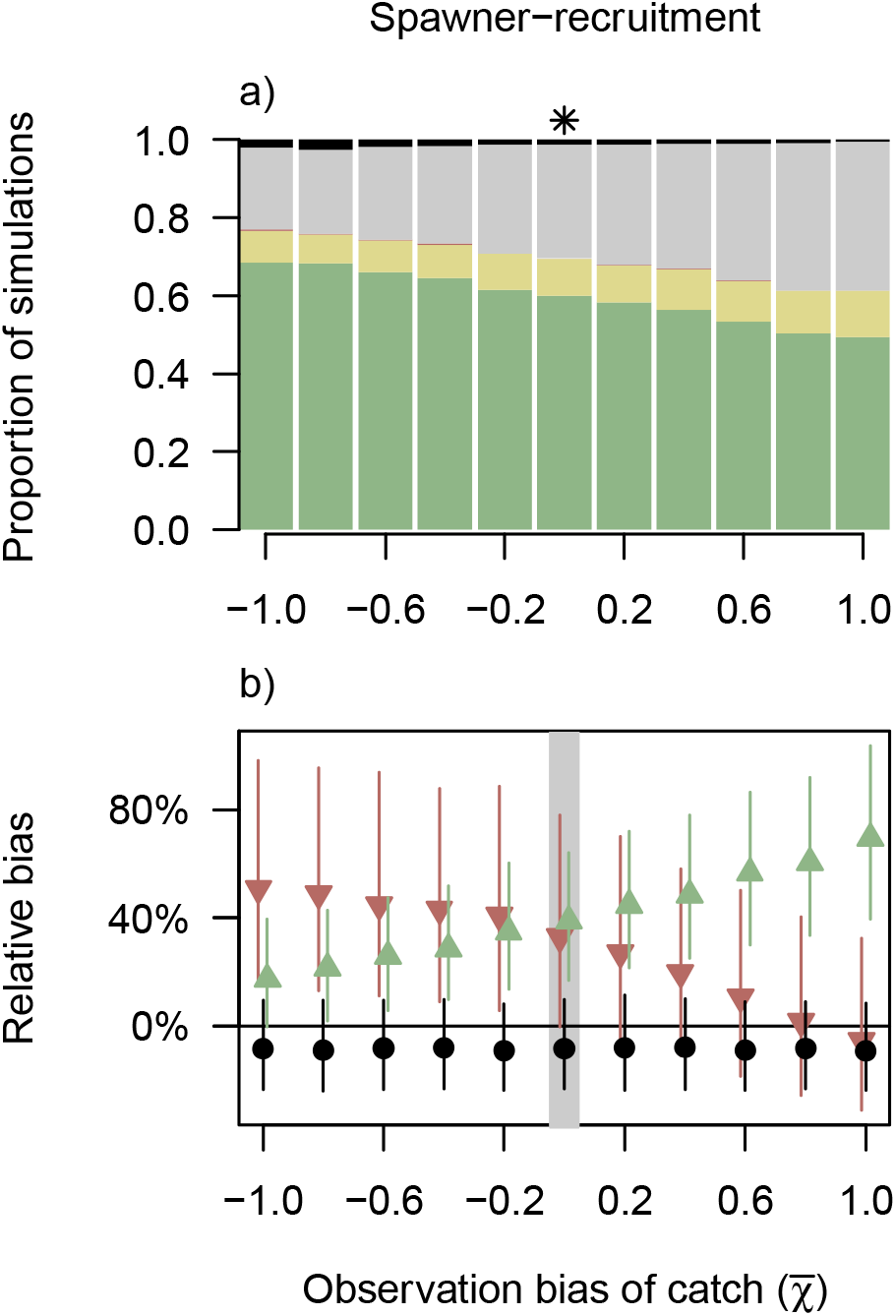
The effect of observation bias in catch (x-axis) on performance measures under the base case. (a) The proportion of simulations with correct green, amber, or red status or pessimistic misclassifications (grey) and optimistic misclassifications (black) under the SR benchmarks. (b) The percent relative bias (median ± interquartile range among 4000 MC simulations) in the current spawner abundance (S_AVG_; black circle) and lower and upper benchmarks (red and green, respectively) under the SR benchmarks. The asterisk in (a) and grey zone in (b) indicate the default parameter value of 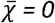. Observation bias in catch ranges from 63% underestimation 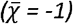 to 271% overestimation 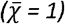. See Online Supplement for results under the low-productivity high-harvest case.

## Discussion

In this study, we quantified the impact of common assumptions in run reconstructions (Figure 2c) on biological status outcomes. In general, the assumptions that we hypothesized might introduce biases (Table 1) had little effect on status outcomes, suggesting that attempts to assess status in the face of limited or uncertain data are worthwhile. In particular, there was almost no effect of declines in monitoring coverage on the accuracy of benchmarks or resulting status outcomes, even in the face of different trends in capacity and reduced synchrony (i.e., zero autocorrelation in recruitment deviates) among spawning populations within the CU. Extreme declines in monitoring may impact assessments – certainly, if no spawning populations are monitored then our ability to assess status will inevitably be compromised – but the current approach to imputing and expanding spawner abundances appeared robust to declines in monitoring coverage in the range documented for the north and central coast (English 2016; Price et al. 2017).

Perhaps unexpectedly, misclassifications were not minimized when the value of Expansion Factor III used in run reconstructions, correcting for observer efficiency, perfectly offset the observation bias in the underlying simulation. The bias in estimated current spawner abundance was lowest when Expansion Factor III matched observation bias, but the positive bias in benchmarks under the base case meant that misclassifications under the spawner-recruitment (SR) benchmarks were minimized when there was no observation bias in spawners. Status outcomes under the percentile benchmarks were unaffected by observation bias, as this bias was assumed to be constant over time and affected the current and historical spawner abundances equally.

Under our base case of high productivity and an abundance-based harvest control rule (HCR) consistent with historical central coast chum salmon harvest rates, most simulations had a true green status, but pessimistic misclassifications as amber were common under both SR and percentile benchmarks. The estimated status from our simulations roughly matched the status outcomes for seven central coast chum CUs from that period, with the majority of CUs having green status under the SR benchmarks and amber status under the percentile benchmarks (Connors et al. 2018). (Note that status of central coast chum salmon CUs has since declined; see the Pacific Salmon Explorer (www.salmonexplorer.ca) for the most up-to-date assessments.) Consistent with the real status assessments, status under the percentile benchmarks tended to be poorer than status under the SR benchmarks. Our simulations attributed this to a higher relative bias in percentile benchmarks, as found for south-coast chum salmon under high-productivity scenarios (Holt et al. 2018). Pessimistic misclassifications may lead to overly conservative management actions, consistent with the Precautionary Principle (Foster et al. 2000), but may also result in foregone harvest and adverse socio-economic consequences (Walters et al. 2019).

For simulations with low productivity and high harvest rates (i.e., true red status), the bias in benchmarks was negative, resulting in a high proportion of optimistic misclassifications. This represents a high-risk management scenario from a conservation standpoint, particularly as the true underlying status is red. Further, the absolute biases in benchmarks were more severe under declines in capacity when status was predominantly red (productivity low) compared to predominantly green (productivity high). The bias in SR benchmarks was particularly sensitive to the underlying true status, presumably because productivity and recruitment, which influence status, also affect the bias in SR parameters (Subbey et al. 2014) that arises due to recruitment-spawner linkage inherent in the data (Walters 1985; Korman et al. 1995) and/or due to error in spawner estimates (Walters and Ludwig 1981; Kehler et al. 2002; Kope 2006). More sophisticated modelling approaches, such as the state-space models that can account for errors in variables and allow for information to be shared among populations and/or CUs, may reduce bias in benchmarks and lead to more robust status assignments (Staton et al. 2020).

Our results suggest that overestimating catch should be avoided. In particular, under low productivity and high harvest rates, optimistic misclassifications associated with overestimating catch and therefore underestimating the lower benchmark, *S*_GEN_, may put populations at further risk. Under the base case of high productivity and an HCR, overestimating catch resulted in more frequent pessimistic misclassifications as the upper benchmark (*S*_MSY_) was overestimated, resulting in CUs with a true green status being estimated as amber. In both cases, the impact of overestimating catch has the potential to significantly bias assessments: overestimating catch by ~50% led to a 5 – 8% increase in misclassification rates. This level of catch overestimation (and higher levels) may occur when fish caught in a Pacific Fisheries Management Area (PFMA) and assigned to the CU that overlaps with that PFMA were actually bound for other CUs. This could occur in mixed-stock fisheries if genetic stock identification is not undertaken to validate assumptions regarding run-timing and migration patterns. Increased efforts to quantify catch composition, run timing, and spatial distribution of Pacific salmon CUs are therefore needed to more accurately estimate harvest rates and minimize misclassifications of biological status.

### Limitations, challenges, and future research

Closed-loop simulation models are powerful tools for evaluating management strategies and quantifying the biases in parameter estimation and status outcomes (Walters 1986; Peterman 2018), but are not without their weaknesses. As with any model, our simulation model was an approximation of reality and thus we had to make a number of assumptions. Both the true population dynamics and assessment submodel assumed a Ricker SR relationship (Ricker 1954), but we recognize that there is considerable model uncertainty. Distinguishing among different models (e.g., Ricker versus Beverton Holt) in assessing status would be challenging, particularly under the declines in capacity that we simulated to capture observed changes in freshwater habitat. The Ricker model is commonly used to set management targets and for simulating population dynamics of Pacific salmon (e.g., Peterman et al. 2000; Peacock and Holt 2012; Fleischman et al. 2013; Holt and Folkes 2015), and a full exploration of other true models was beyond our scope.

We did not implement a bias correction when simulating log-normal recruitment deviates, consistent with other Pacific-salmon simulation studies (e.g., Dorner et al. 2013; Cunningham et al. 2019). This introduced a positive bias in mean recruitment and subsequent estimates of productivity. Alternatively, one could include a bias correction when simulating recruitment by assuming the mean of *ν_y,j_* in eq. (7) is 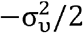, and this may be best practice moving forward (e.g., Hicks et al. 2020). However, there is no clear consensus about whether this correction is necessary when parameters are estimated from simulated data in an assessment model without a 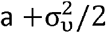 adjustment (Hilborn and Walters 1992).

We considered true SR dynamics to operate at the spatial scale of spawning populations (i.e., individual streams), but there is evidence that the processes influencing productivity and density-dependence may operate at broader, regional spatial scales (e.g., Malick and Cox 2016). Other simulation models have incorporated straying among spawning populations within a CU (e.g., Peacock and Holt 2012; Holt and Folkes 2015). While that approach incorporates density-dependence that may occur at broader spatial scales, it also requires additional assumptions to be made about the probabilities of straying among streams, which are largely unknown.

Simulating true dynamics at the scale of individual spawning populations also complicates the calculation of true status at the CU-level from SR benchmarks. We chose to calculate SR benchmarks at the spawning-population-level and then sum across spawning populations to calculate *S*_MSY_ and *S*_GEN_ at the CU-level. There are other approaches to calculate aggregate benchmarks, but each has its own potential biases. For example, SR relationships could be fit to the “true” data aggregated at the CU-level and SR benchmarks calculated from the resulting CU-level estimates of productivity and density dependence. The way in which spawning-population-level benchmarks are aggregated to CU-level benchmarks may affect performance in our simulations, and a full exploration of how different methods of aggregation affect our results warrants future consideration.

Despite these limitations, the simulation model that we adapted and applied is flexible enough that it can accommodate different Pacific salmon species and life-history traits, opening the door to future work investigating the impact of different assumptions and the impact of the assumptions that we focused on under additional scenarios. Here, we considered a relatively simple run-reconstruction model, but further work is needed to quantify how observation errors and uncertainty in the spatial and temporal distribution of returns affects status outcomes when more complex run-reconstruction models are used. Temporal shifts in biological parameters, including age-at-return (e.g., associated with environmental change and selective fisheries) and productivity (non-stationarity has been widely observed in Pacific salmon; e.g., Peterman and Dorner 2012; Malick and Cox 2016; Dorner et al. 2017), are also areas that warrant further exploration. Additional simulations could also help inform methods in the assessment process, such as the optimal time-series length for detecting changes, whether benchmarks should be updated with each assessment, and the best analytical approach to calculate SR benchmarks (e.g., Bayesian hierarchical models vs. single-stock ordinary least squares).

### Conclusions

Pacific salmon are one of the most data-rich groups of fish due to their high economic, social, and cultural value, but nonetheless our knowledge of their dynamics is uncertain. Assessing the biological status of Pacific salmon CUs is a conservation and management priority given the continued declines of many stocks (e.g., COSEWIC 2016, 2017) and escalating threats to salmon conservation. Current government-led integrated status assessments under the WSP include expert opinion on the data uncertainties and risks unique to each CU (e.g., DFO 2015, 2016, 2018a), making the process time and resource intensive. This has limited their timely application to all 460+ Pacific salmon CUs in Canada (DFO 2019), and also means that the process is not entirely transparent or reproducible. The Pacific Salmon Foundation has implemented data-driven biological status assessments based on a subset of the indicators and benchmarks recommended under the WSP, with the results accessible through the Pacific Salmon Explorer (www.salmonexplorer.ca). These data-driven assessments, similar to those undertaken by other management and conservation organizations (e.g., Marine Stewardship Council, COSEWIC, Pacific Salmon Commission), require assumptions to estimate spawner and recruit time series for CUs. Given the lack of expert scrutiny of the data for each individual CU under this data-driven approach, it was important to understand how these assumptions may bias status outcomes under a range of biological and management scenarios.

We found that the data-driven biological status assessments were relatively insensitive to common assumptions in expanding spawner abundances within the parameter ranges we explored, but the rate and type of status misclassifications depended on the underlying status of the CU and may be of greater concern for CUs with poor status. To ensure the accuracy of data-driven status assessments, increased efforts to collect data on catch composition, age-at-return, and spawner abundances are needed. Such information will help, for example, to define plausible ranges of error in catch estimation to lend confidence to estimates of recruitment and thus assessments under SR benchmarks. Nonetheless, our research suggests that current efforts to assess status in the face of imperfect and incomplete data are worthwhile for central coast chum salmon and other similar Pacific salmon populations and can provide a timely approach to assessing status for CUs that complements more thorough integrated status assessments.

## Acknowledgements

The authors thank the Pacific Salmon Foundation’s Population Science Advisory Committee, in particular David Peacock and Randall Peterman, and Steven Cox-Rogers and Charmaine Carr-Harris for their valuable feedback throughout this project. Thanks to Doug Stewart and Kate McGivney for sharing their insights on central coast chum salmon and Brooke Davis and two anonymous reviewers for feedback on earlier drafts. This research was funded by the Pacific Salmon Foundation’s Salmon Watersheds Program.

## Appendices

## Appendix A: Table of default parameter values

**Table A1.**
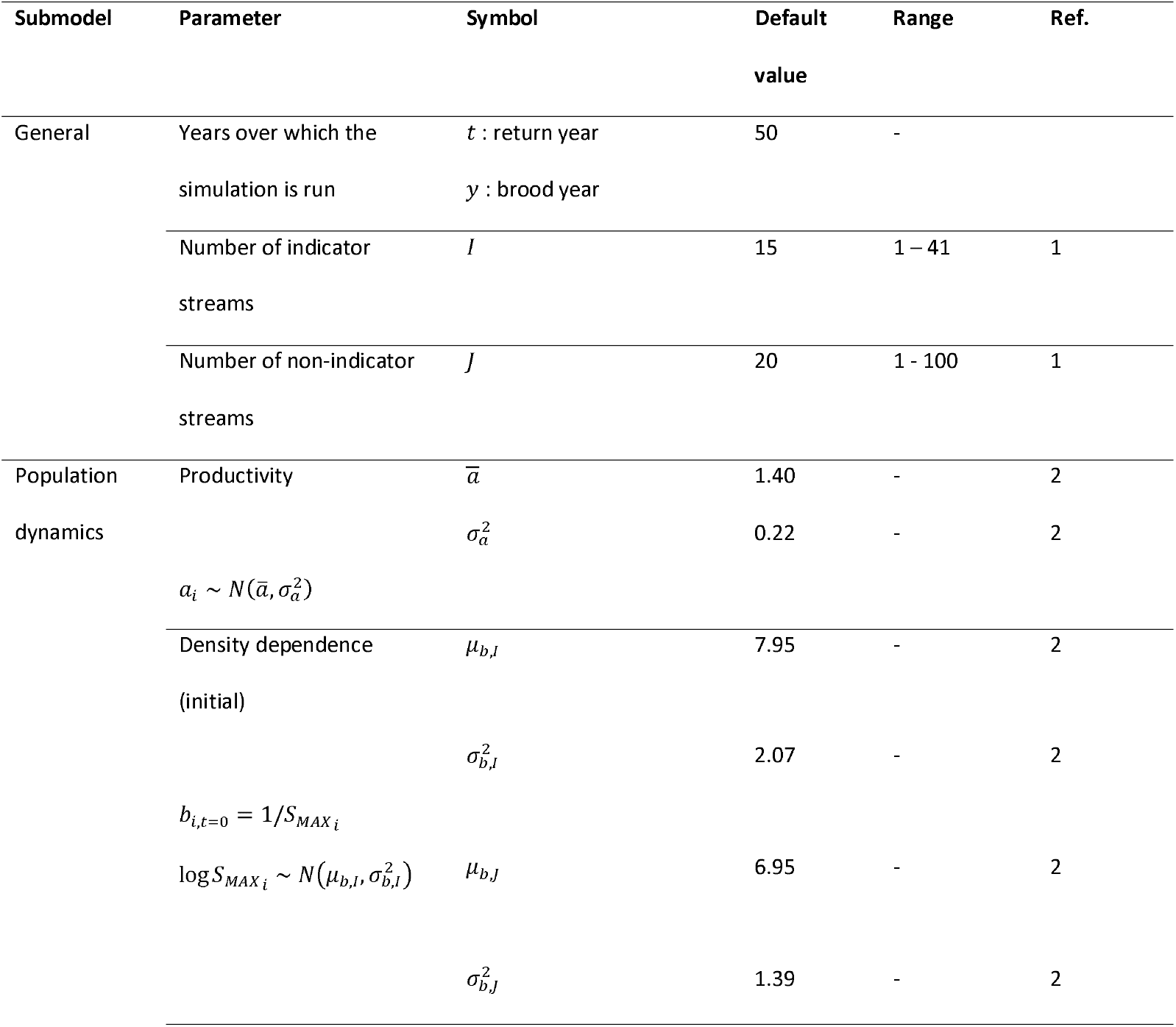
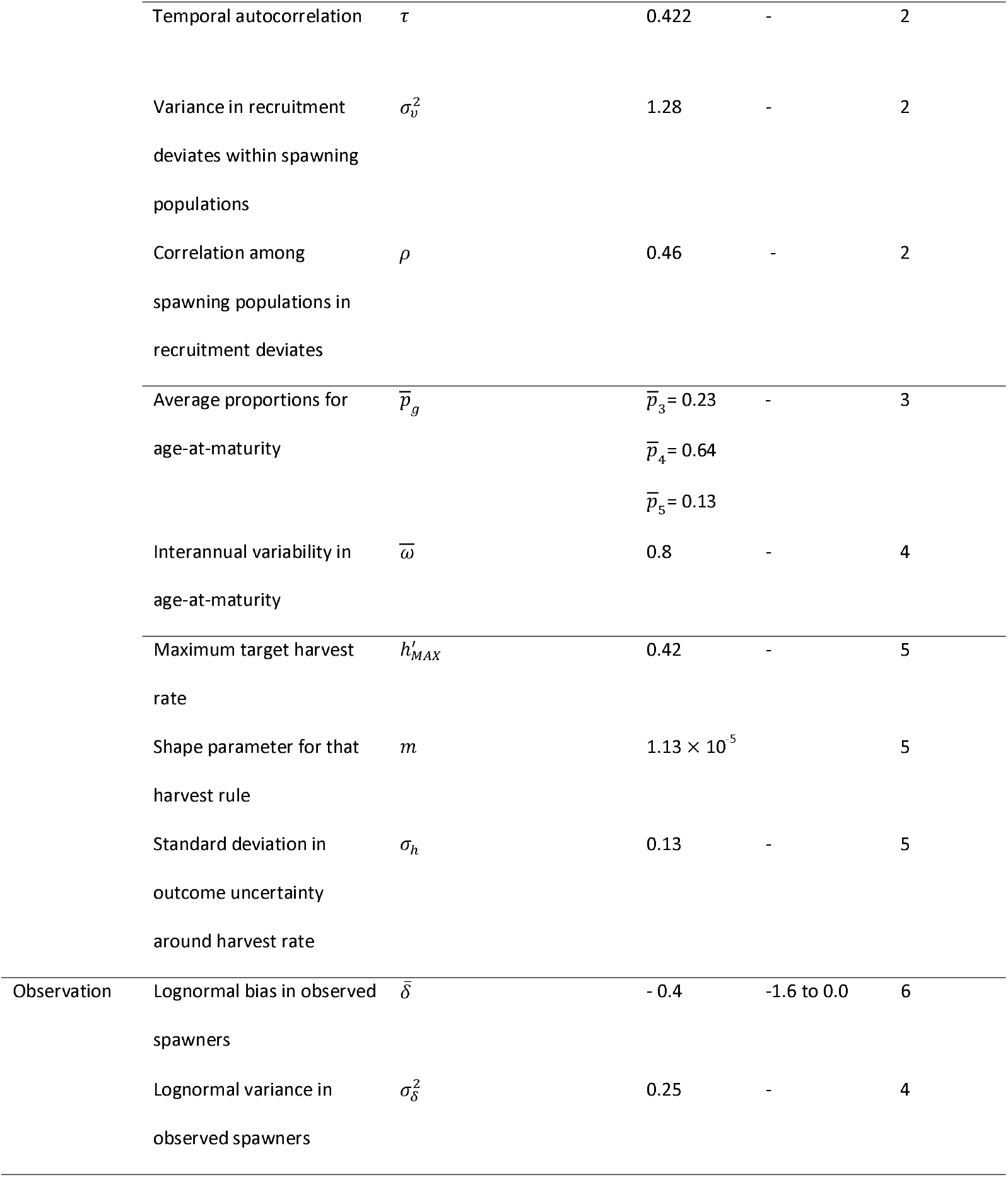
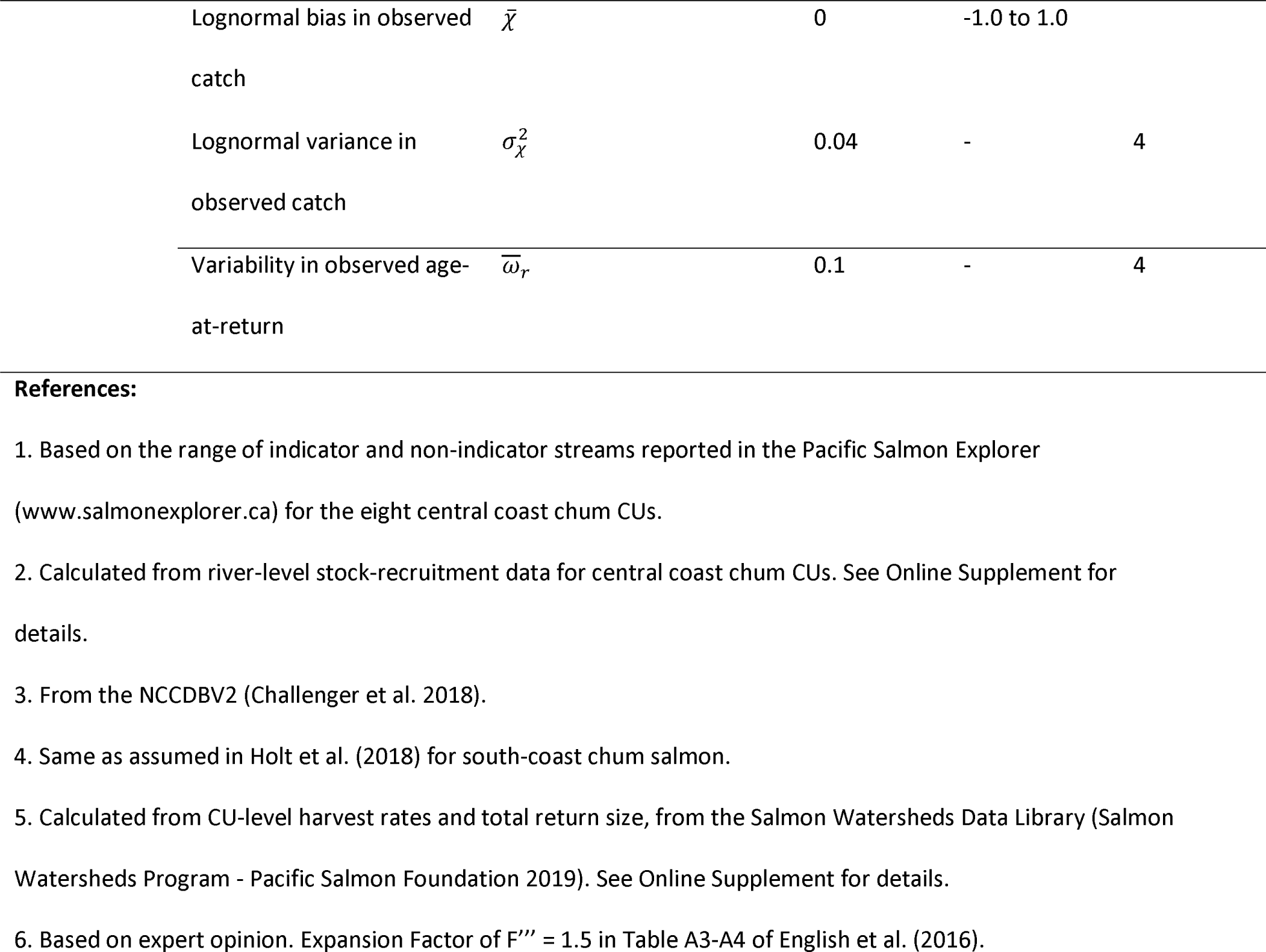
Default values for parameters in the simulation model that were used unless otherwise specified (e.g., in sensitivity analyses). See text for further explanation of the values and the Online Supplement for details of estimation for those based on raw data. For parameters that were part of sensitivity analyses, the range in parameter values that was explored is highlighted.

