## Supplementary material for "Evaluating the consequences of common assumptions in run reconstructions on Pacific-salmon biological status assessments": Online Supplement

Eric Hertz, Carrie A. Holt, Brendan Connors, Cameron Freshwater, and Katrina Connors

### 1 Supplemental methods

The code for the model and results is available at <https://github.com/sjpeacock/run-reconst-sim_PSF>. Version 1.0, used in this paper, is archived at [doi:10.5281/zenodo.3971276](https://zenodo.org/badge/latestdoi/161822916) and was implemented in R version 4.0.2.

#### 1.1 Expansion factors

**Expansion Factor I,** $F'_{t/d}$, expands the observed spawner abundances in indicator streams to account for indicator streams that are not monitored in a given year. It is calculated for each year $t$ within decade $d$ of the spawner time series, and relies on a decadal contribution of each indicator stream to the total escapement to all indicator streams, $P_{d,i}$ (English et al. 2016). The calculation of this decadal contribution requires at least one estimate from each indicator stream for the decade. If a decade does not contain sufficient information (i.e., one or more indicator streams are not monitored at all in a decade), then a reference decade is used to calculate $P_{d,i}$. This reference decade is chosen to be: (1) the closest decade (historical or future) with sufficient information, or failing (1), (2) the 20-year period from 1980-1999 (Challenger et al. 2018).

For each decade (or reference decade if insufficient information) $d$, the average number of spawners returning to indicator stream $i$ in decade $d$ is calculated as:

$$\overline{S}_{d,i}=\frac{\sum_{t=1}^{Y_{d,i}} {\overset{^}{S}}_{t,d,i}}{Y_{d,i}}$$

(S1)

where $Y_{d,i}$ is the number of years for which spawner estimates are available within decade $d$ for stream $i$. From the average number of spawners for all indicator streams, the decadal proportional contribution of each indicator stream is calculated as:

$$P_{d,i}=\frac{\overline{S}_{d,i}}{\sum_{i=1}^{I} \overline{S}_{d,i}}$$

(S2)

where $I$ is the total number of indicator streams.

Expansion Factor I is then calculated for each year, $t$, within the decade, $d$, based on the decadal contributions and which streams were monitored or not in a given year:

$$F'_{t,d}=\left( \sum_{i=1}^{I} \left\{ P_{d,i} w_{t,d,i} \right\} \right)^{-1}$$

(S3)

where $w_{t,d,i}$ is 1 if stream $i$ was monitored in year $t$ and 0 if stream $i$ was not monitored in year $t$. The expansion factor $F'_{t,d}$ is then multiplied by the sum of the observed spawners in all indicator streams to yield the expanded estimate of spawner abundances in all indicator streams for the CU:

$$S'_{t,d}=F'_{t,d}\sum_{i=1}^{I} {\overset{^}{S}}_{t,i}.$$

(S4)

**Expansion Factor II,** $F{''}_{d}$ expands the escapement to all indicator streams, $S'_{t}$, to account for non-indcator streams. Unlike Expansion Factor I, this is calculated for each decade (rather than each year) and then applied to all years within a decade. Like Expansion Factor I, there needs to be sufficient information within the given decade in order to calculate $F{''}_{d}$, or else a reference decade is chosen. To date, these reference decades have been selected based on expert opinion. In our simulation framework, we had to define the selection of reference decades based on quantitative criteria (Figure S1).

1. Every indicator stream is monitored at least once within a decade

a. The number of non-indicator streams monitored at least once varies <=10% across decades

Use current decade as reference decade

b. The number of non-indicator streams monitored at least once is significantly higher (>10%) in one decade comnpared to any other decade

Use decade with the best coverage of non-indicator streams as reference decade

c. The number of non-indicator streams monitored at least once is >10% higher in two or more deacdes compared to other decades

For decades when the number of non-indicator streams monitred at least once is signficiantly lower, use the closest historical or future decade with higher coverage

2. Not every indicator stream is monitored at least once in every decade

a. The number of non-indicator streams monitored at least once varies <=10% across decades

Use the reference decades for Expansion Factor I, which rely on coverage of indicator streams only.

b. The number of non-indicator streams monitored at least once varies >10% across decades

Calculate the hypothetical reference decade for non-indicators based on the criteria for Expansion Factor 1.

(i) If the reference decades for indiciator and non-indicator streams are the same, then use those.

(ii) If not, use the period from 1980-1999 or, if the number of non-indicator streams is significantly different from 1980-1989 and 1990-1999, then use the entire timeseries.

Figure S1. Our flow chart for selecting a reference decade when calculating Expansion Factor 2.

Expansion Factor II is calculated as:

$$F_{d}^{''}=\frac{\sum_{i=1}^{I} \overline{S}_{d,i}+\sum_{j=1}^{J} \overline{S}_{d,j}}{\sum_{i=1}^{I} \overline{S}_{d,i}}$$

(S5)

where $\overline{S}_{d,i}$ and $\overline{S}_{d,j}$ are the decadal average number of spawners in indicator and non-indicator streams, respectively, calculated by eqn. (S1). $J$ is the total number of populations in non-indicator streams. The adjusted total number of spawners in both indicator and non-indicator streams is then calculated as:

$$S_{t,d}^{''}=F_{d}^{''} S_{t,d}^{'}$$

(S6)

Finally, the number of spawners in both indicator and non-indicator streams, $S_{t,d}^{''}$, is multiplied by **Expansion Factor III** to account for streams that are never monitored and for observer (in)efficiency. Expansion Factor III is determined by the regional DFO staff familiar with the escapement monitoring techniques used in each statistical area and is given as $F^{‴}$ = 1.50 for all north and central coast chum CUs in Table A3 and A4 of English et al. (2016). The final expanded number of spawners (i.e., escapement) to the CU for year $t$ is given by $S_{t}^{‴}=F^{‴}S_{t,d}^{''}$.

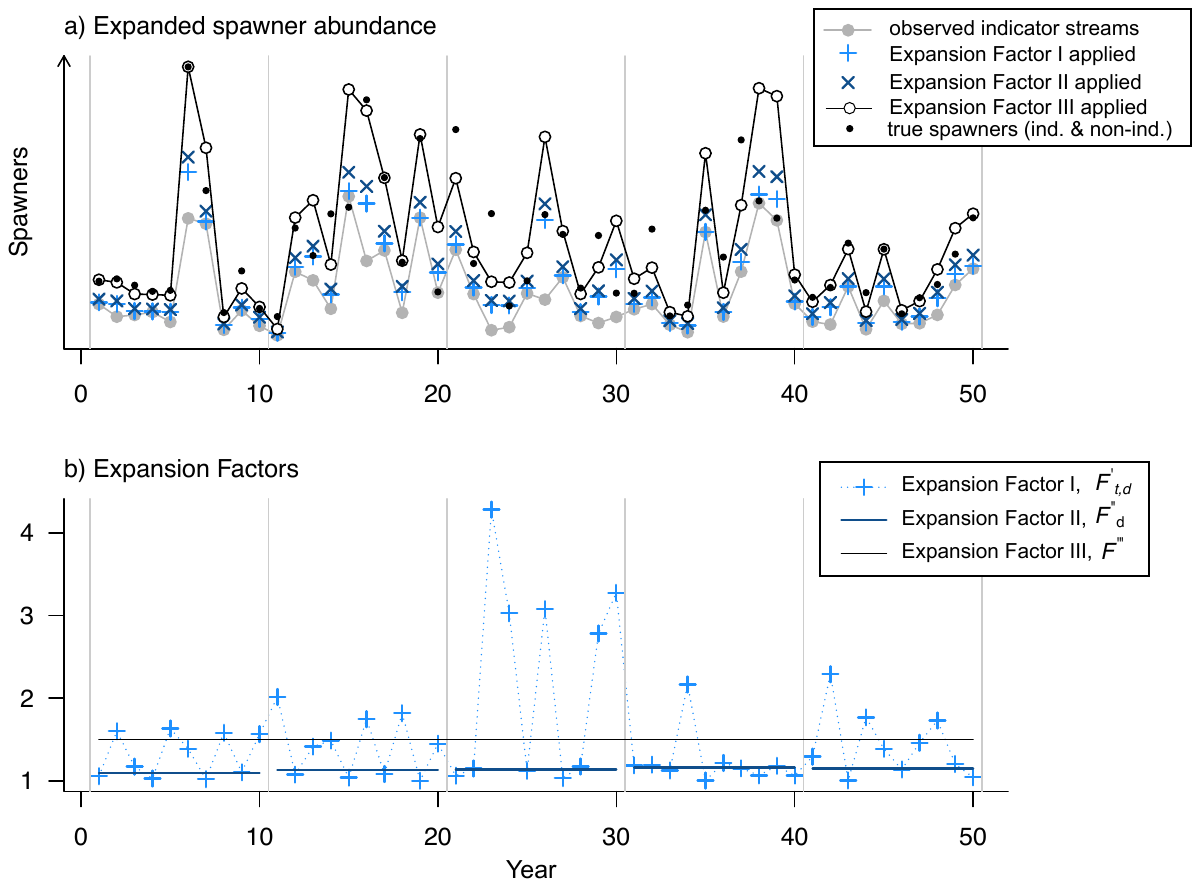

Figure S2. a) Time series of observed spawner abundance to indicator streams (grey points and line), the imputed spawner abundance to indicator streams after the application of Expansion Factor I (light blue +), the expanded spawner abundance to both indicator and non-indicator streams after the application of Expansion Factor II (dark blue x), and the final expanded spawner abundance after application of Expansion Factor III (black open points and line). The small closed points are the true spawner abundance. B) The values of Expansion Factor I (F’_t_), which is calculated annually (light blue + and dotted line), Expansion Factor II (F’’d) which is calculated for each decade (horizontal dark blue segments), and Expansion Factor III (F’’’) which is constant over time (horizontal black line). The values shown here are the Expansion Factors applied in (a).

#### 1.2 Performance submodel

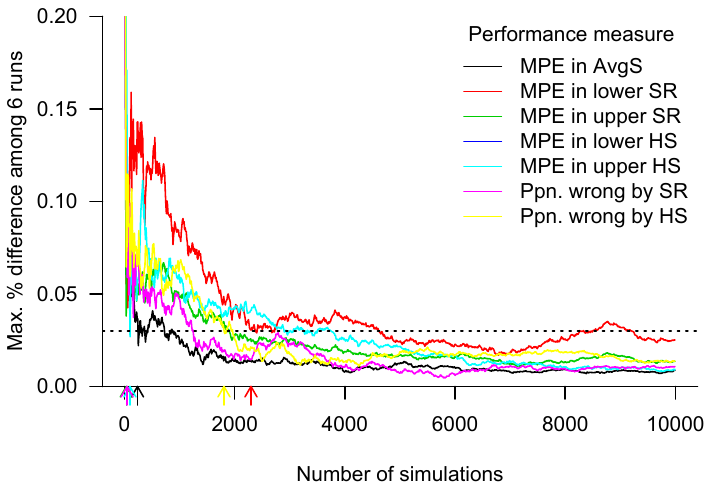

Figure S3. The maximum percent difference between the seven performance measures (relative bias for average spawners and four benchmarks and the proportion of simulations with incorrect status assignment by the spawner-recruitment and percentile metrics) over six independent runs of 10,000 MC simulations each. The maximum percent difference was calculated as the largest value of the performance measure minus the smallest value divided by the largest value. The horizontal dotted line is at 0.03, or 3%, which was our threshold for sufficient number of MC trials. We chose 4000 simulations for all of our subsequent analyses, which was sufficient to ensure the difference among simulations was < 3% across performance measures.

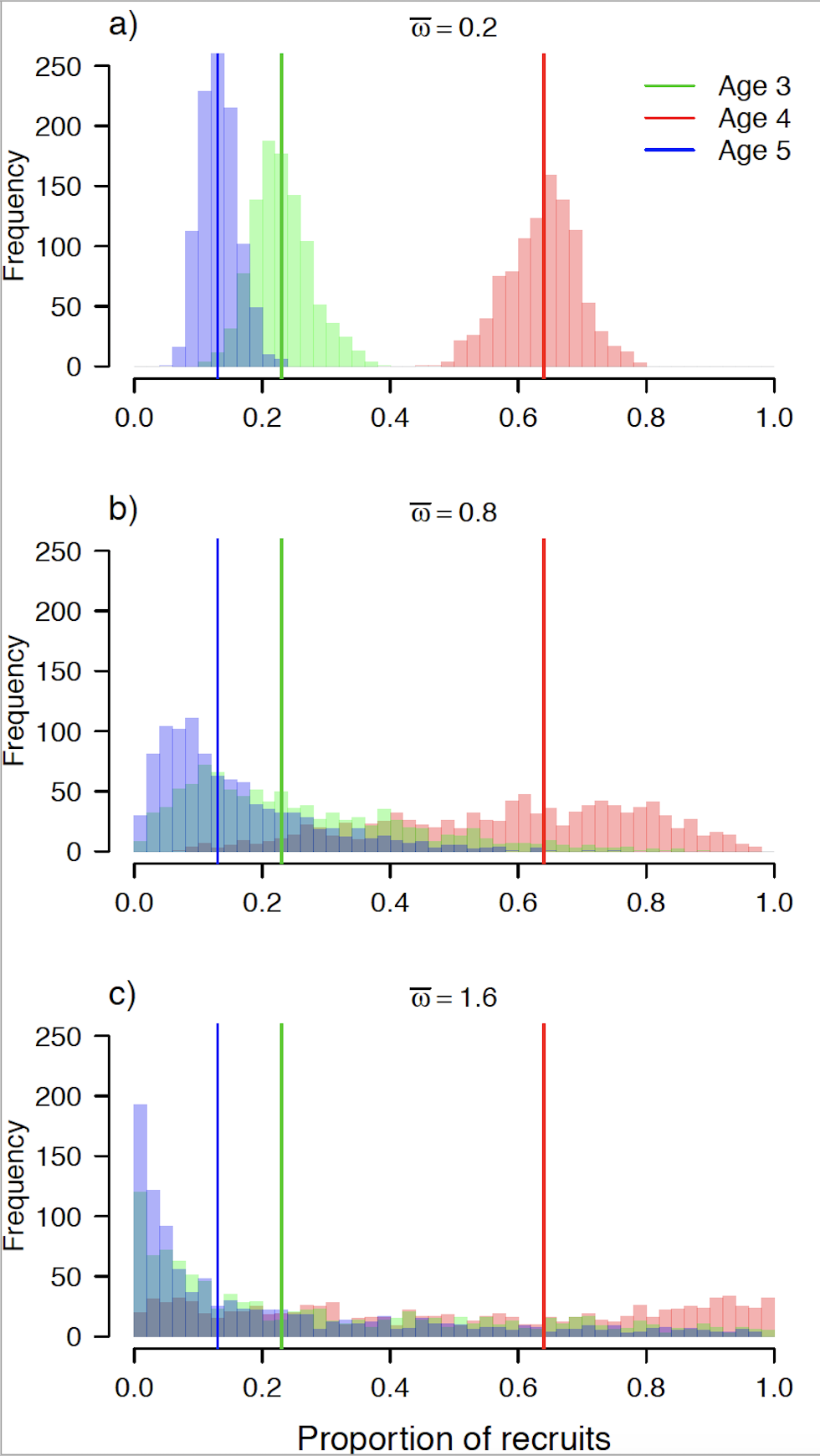

Figure S4. An example of how the parameter $\bar{\omega}$ affects the distribution of age-at-maturity from which annual proportions are drawn (eq. (2) in main text). The histograms show 1000 random draws of ages-at-maturity from eq. (2) using mean proportions at age of $\bar{p_{3}}$ = 0.23 (green)**,** $\bar{p_{4}}$ = 0.64 (red), and $\bar{p_{5}}$ = 0.13 (blue), and three different levels of interannual variability: a)  $\bar{\omega}$ = 0.2, b)  $\bar{\omega}$ = 0.8 (default value), and c) $\bar{\omega}$ = 1.6.

### 2 Parameter calculations

#### 2.1 Target harvest rates

We fit eq. (4) from the main text, describing the target harvest rate as a function of total return size (Holt and Peterman 2008), to CU-level data of estimated harvest rates at total return size for five central coast chum CUs (Salmon Watersheds Program - Pacific Salmon Foundation 2019). Two CUs (Rivers Inlet and Smith Inlet) were excluded despite having data because harvest rates have been zero in those CUs over recent decades. One other CU (Bella Coola River-Late) lacked data on total run size. We fit the model in R (R Development Core Team 2018) using the function nls. The least-squares estimates for parameters were: $h_{MAX}^{'}$ = 0.42 and $m$ = 1.13 $\times$ 10^-5^. We also considered a model with an additional parameter for a “protection level”, which is the return below which harvest rates are zero (see Holt and Peterman 2008). However, this parameter was estimated to be zero and also did not make sense given that we applied a minimum harvest rate of $h_{t}^{'}=$0.05 regardless of return size.

#### 2.2 Ricker parameters

We fit a simple linearized Ricker model to spawner-recruit pairs for spawning populations in 181 streams (henceforth “populations”) from eight central coast chum CUs with individual productivity and density-dependence parameters for each population:

$$\log_{e} \left( \frac{R_{i,t}}{S_{i,t}} \right)=a_{i}-b_{i}S_{i,t}+\epsilon_{i,t}$$

(S7)

We fit eqn. (S7) in R (R Development Core Team 2018) using the package lme4 (Bates et al. 2015), including a random effect of CU on the intercept (i.e., productivity). There was a total of 5724 spawner-recruitment pairs from 181 streams, spanning brood years 1954-2010.

From these model fits, we calculated

1) mean ($\bar{a}$) and variance ($\sigma_{a}^{2}$) in productivity among all 181 populations,
2) mean and variance in capacity, where capacity was calculated as -log(*b*) = log(*S*_MAX_). The mean and variance in capacity were calculated separately for indicator and non-indicator streams.
3) the residual variance within populations,
4) the average correlation in residuals between each pair of populations (n = 15 985 pairs, excluding pairs for which a correlation could not be calculated because they did not have spawner-recruit data in overlapping years), and
5) the temporal autocorrelation in residuals within populations.

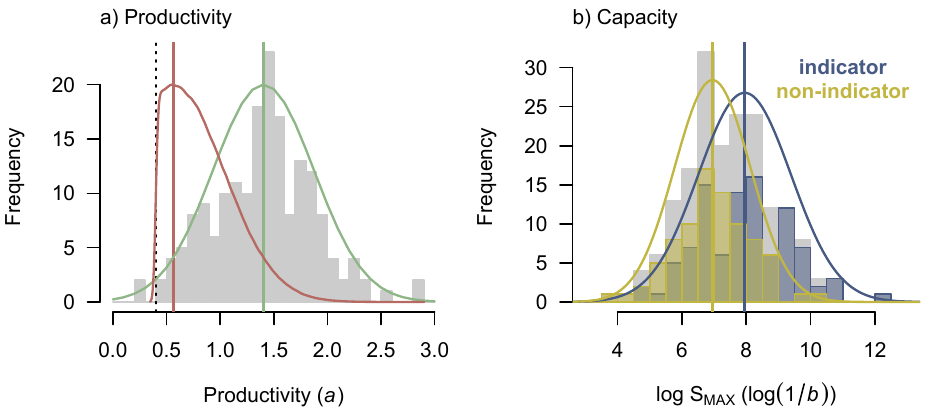

Figure S5. Ricker productivity and density-dependence (capacity) parameters estimated from spawner-recruitment data from spawning populations in 181 streams over eight central coast chum CUs. a) Productivity mean was 1.4 (vertical green line) and variance $\sigma_{a}^{2}$ = 0.22, resulting in a distribution shaped like the green curve from which parameters were drawn for our high-productivity base case. For low-productivity cases, we set the mean to the 2.5^th^ percentile of the estimated productivity parameters (vertical red line). When incorporating our constraint that a > 0.4, this led to a distribution shaped like the red curve. b) Density-dependence parameters differed between indicator and non-indicator streams, with indicator streams having on average higher capacity. Capacity was log-normally distributed, and is shown here as log(S_MAX_), which is equal to -log(b). The blue and gold lines show the distributions from which S_MAX_ for indicator and non-indicator streams were drawn in simulations. The underlying grey histogram shows the capacity parameters for both indicator and non-indicator streams combined.

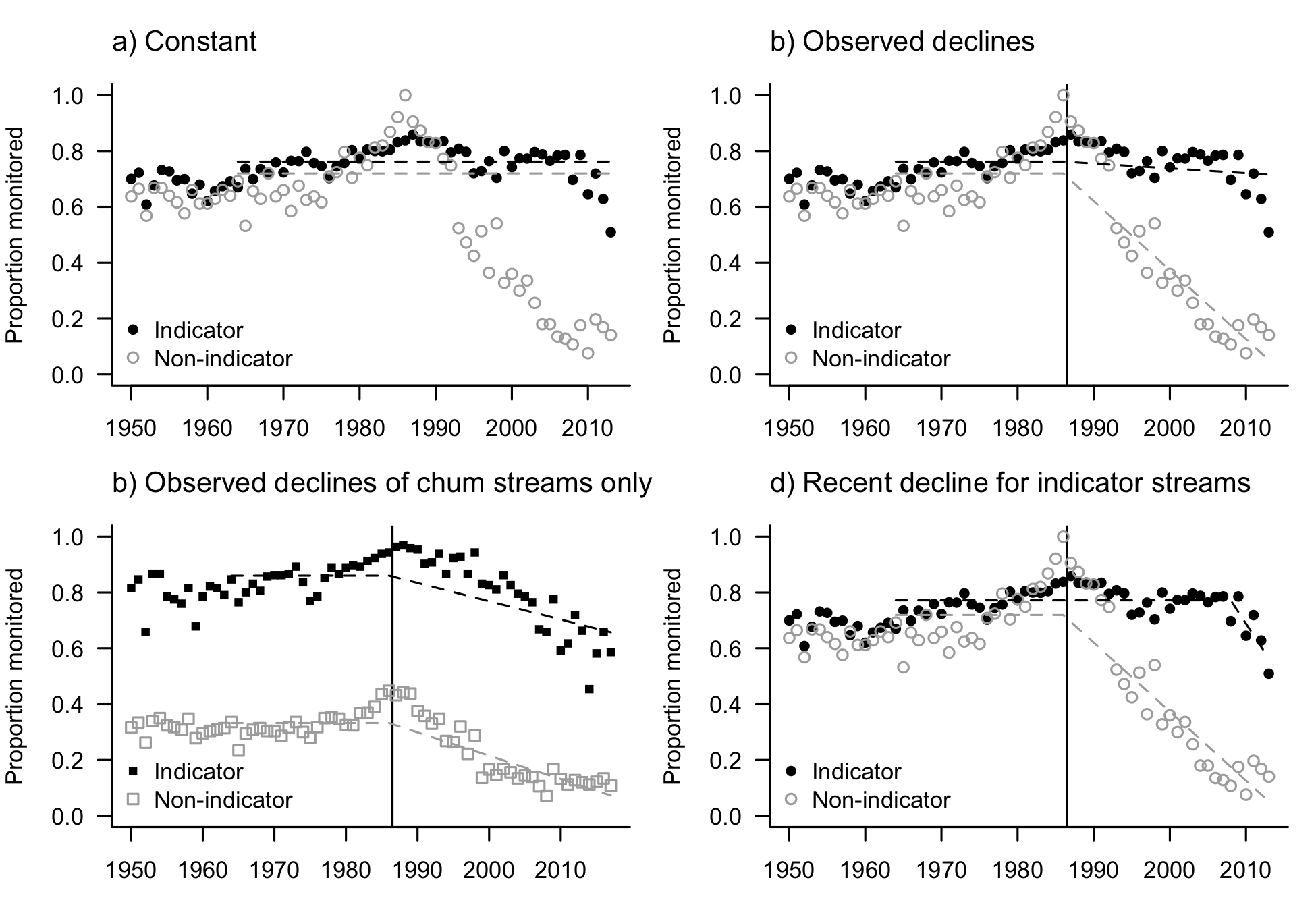

Figure S6. Empirical data on proportion of indicator and non-indicator streams monitored for all Pacific salmon systems in north and central BC (circles; a,b,d) and just central coast chum salmon systems (squares; c) from English et al. (2016). We investigated the impact of changes in monitoring coverage under four scenarios (dashed lines): (a) constant monitoring coverage at pre-1980s levels (0.762 and 0.719 for indicator and non-indicator, respectively); (b) a linear decline from historical monitoring coverage since the 1985 of -0.047 and -0.667 for indicator and non-indicator, respectively; (c) a linear decline from historical monitoring coverage of central coast chum streams (0.86 and 0.33) of -0.20 and -0.24 for indicator and non-indicator, respectively; and (d) a recent (over the last 5-years) declines across all indicator systems from 0.77 by -0.21 (non-indicator kept as in b with declines since the mid-1980s). In the main text, we presented results from a and b, and in the Online Supplement we present results from scenarios c and d. Note that the data shown in a, b, and d are the same, while c shows monitoring coverage chum salmon streams only.

### 3 Supplemental results

Here, we present additional results under default parameterization and results of sensitivity analyses under all three cases. As described in the main text, the first case (“base case”) simulated high productivity with a conservative abundance-based harvest control rule (HCR). The second case (not discussed in the main text) was intermediate with low productivity and constant moderate target harvest rate of $h_{t}^{'}=0.42$. The third case was low productivity and constant high target harvest rate of $h_{t}^{'}=0.60$.

We also include results of two additional sensitivity analyses focused on biological parameters in the true population dynamics. We investigated the sensitivity to interannual variability in age-at-return (Fig. S25) because it was hypothesized to affect the validity of assumptions about age-at-return applied in run reconstructions (namely that an average age-at-return can be applied across all years).

Second, we investigated the sensitivity to the standard deviation in the recruitment deviates (Fig. S26-S27) because our empirical estimation of recruitment variability did not account for observation error and thus was likely an overestimate. We found that spawner-recruitment (SR) benchmarks were more sensitive to changes in the standard deviation in recruitment deviates ($\sigma_{\upsilon}$) than were percentile benchmarks because more variable recruitment increased the contrast in spawner-recruitment data and, thus, affected estimates from the Ricker model that were used in calculating SR benchmarks. As with most of our other results, the effect of changing $\sigma_{\upsilon}$ depended on the underlying status (i.e., productivity and harvest). When true status was mostly green (high productivity/HCR), pessimistic misclassifications increased significantly with increases in $\sigma_{\upsilon}$ (Fig. S26a), due to overestimation of both upper and lower benchmarks (Fig. S26c). When true status was mixed (low productivity/moderate harvest), the rate of pessimistic misclassifications increased (as in the base case) but the rate of optimistic misclassifications declined (Fig 26e). This can be explained by a decline in the proportion of simulations with true amber status with increasing $\sigma_{\upsilon}$ (Fig. 27b) combined with increases in the upper benchmark (Fig. 26g). When true status was mostly red (low productivity/high harvest), the results and mechanism were similar to the second case: optimistic misclassifications declined with $\sigma_{\upsilon}$ (Fig. 26i) due to increasingly positive bias in benchmarks (Fig. 26k) combined with fewer simulations having a true red status (Fig. 27c).

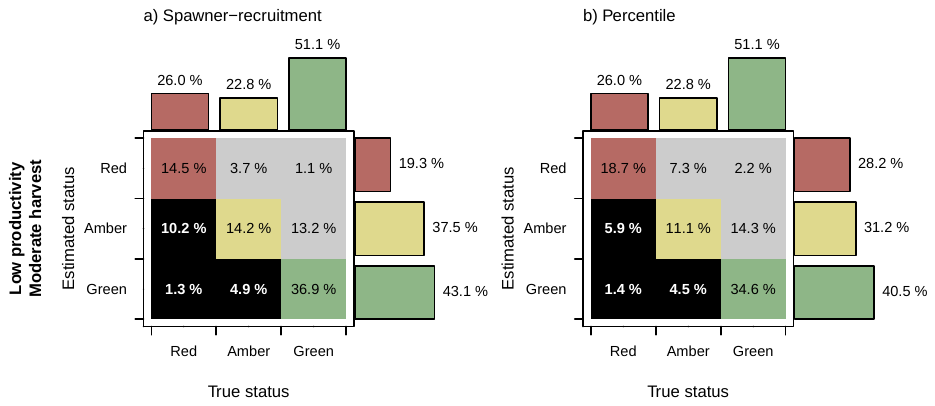

Figure S7. Estimated status according to the spawner-recruitment benchmarks (a) and the percentile benchmarks (b), over the true status for a low-productivity moderate-harvest case (not in main text). The grey zone indicates pessimistic misclassifications, which may lead to overly conservative management actions, and the black region indicates optimistic misclassifications, which may lead to overly risky management actions. See Figure 6 of main text for corresponding results under high-productivity and an HCR (base case) and low-productivity high-harvest.

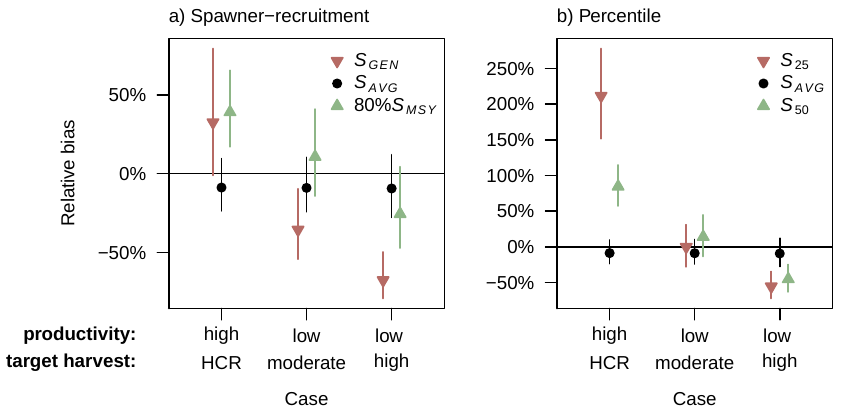

Figure S8. Relative bias in a) spawner-recruitment benchmarks, b) percentile benchmarks, and current spawner abundance (median $\pm$ interquartile range among 4000 MC simulations) under three cases (x-axis): (1) high productivity and a harvest control rule (HCR) and (2) low-productivity moderate-harvest, and (3) low-productivity high-harvest. See main text for results from cases (1) & (2) and Figure S6, above, for case (3).

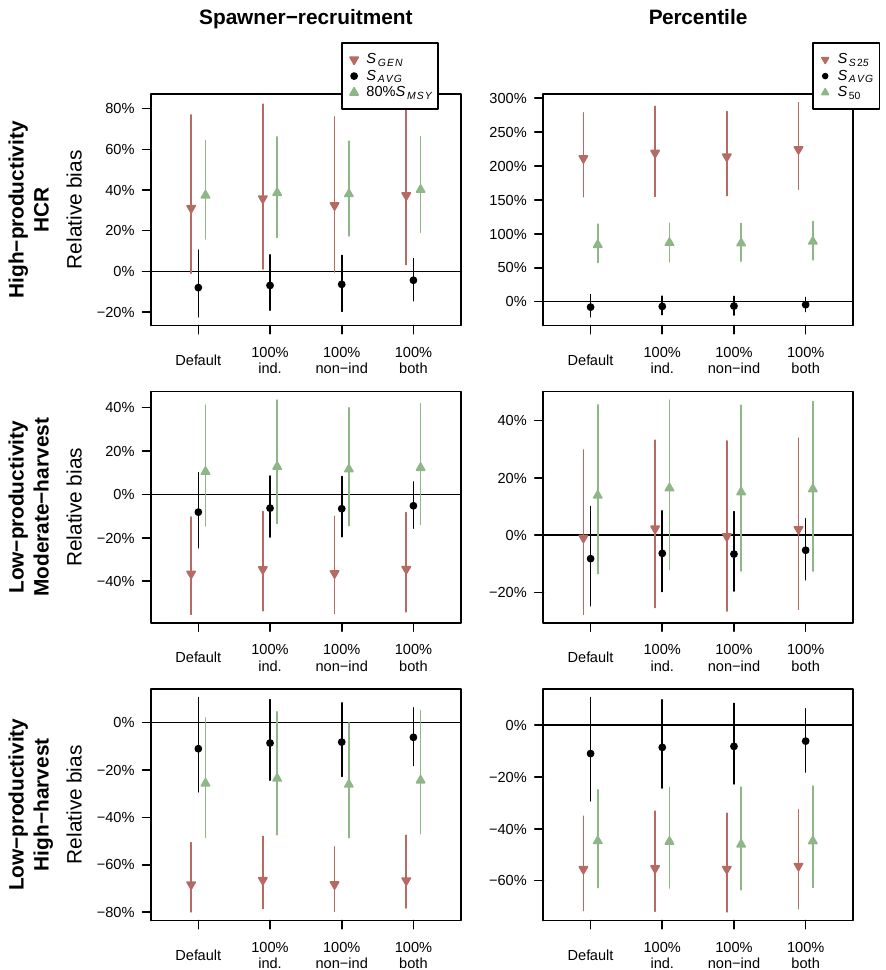

Figure S9. Relative bias in spawner-recruitment benchmarks (left), percentile benchmarks (right), and current spawner abundance (median $\pm$ interquartile range among 4000 MC simulations)(see Figure S8 for legend). The three rows depict three cases for productivity and harvest, and the x-axis in each panel shows the default decline in monitoring coverage, with 100% monitoring coverage of indicator streams (Expansion Factor I = 1.0), 100% monitoring coverage of non-indicator streams (Expansion Factor II = 1.0), and 100% monitoring of both indicator and non-indicator streams (Expansion Factors I and II not required).

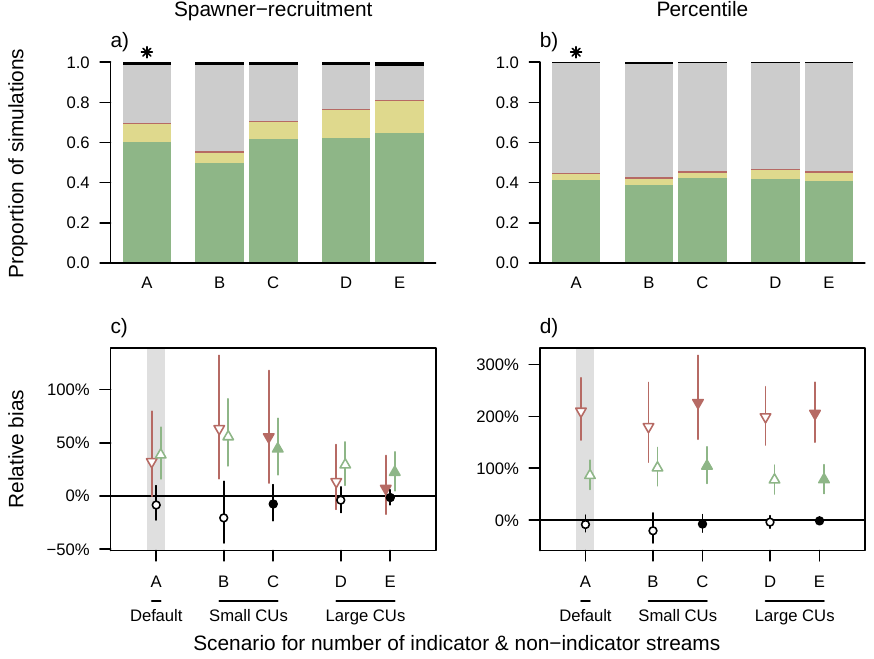

Figure S10. Effect of CU size and proportion of populations that were indicator streams under the base case of high productivity/HCR. The performance measures under five different scenarios for the size of CUs (A = 35 populations; B,C = 10 populations; D,E = 140 populations) and proportion of populations that are indicator streams (A = 75%; B,D = 30% (open points); C,E = 80% (closed points)). a-b) The proportion of simulations with correct green, amber, or red status, pessimistic misclassifications (grey) and optimistic misclassifications (black). c-d) The relative bias in the estimated current spawner abundance (black circle) and the lower (red down arrow) and upper (green up arrow) benchmarks for both metrics. Asterisk in panels (a) and (b) denote the default parameterization (“A”).

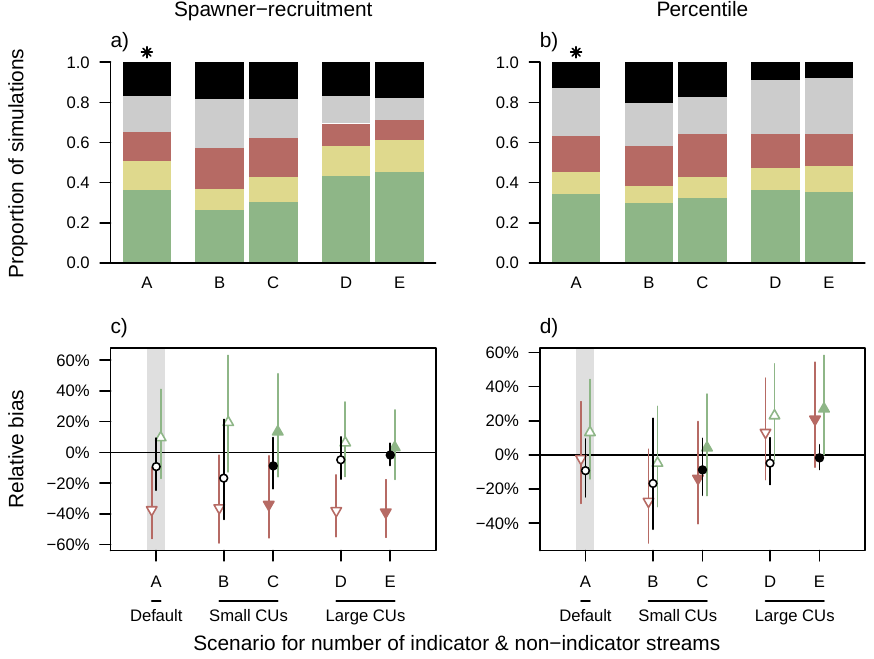

Figure S11. As for Figure S10, but for the low-productivity moderate-harvest case.

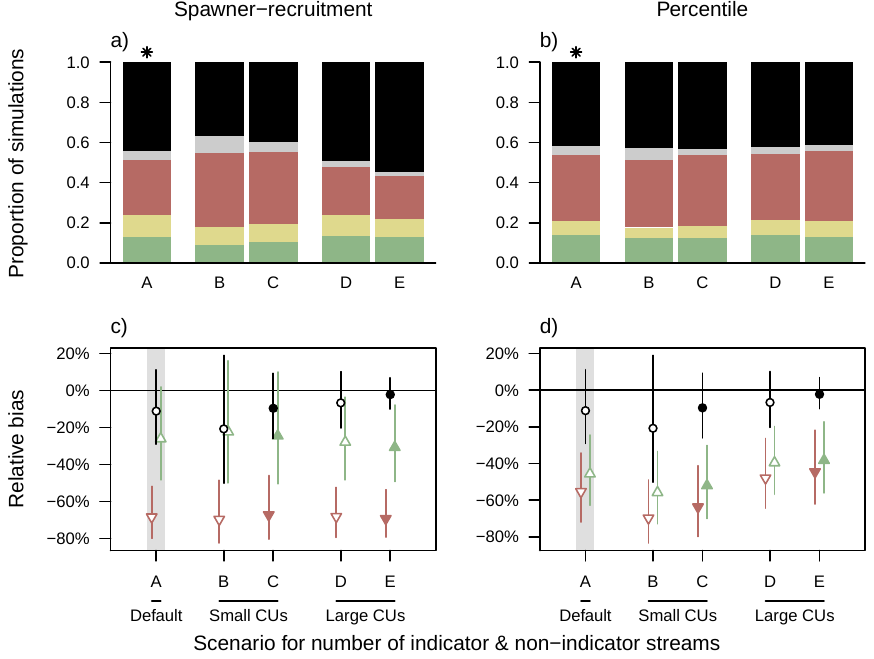

Figure S12. As for Figure S10, but for the low-productivity high-harvest case.

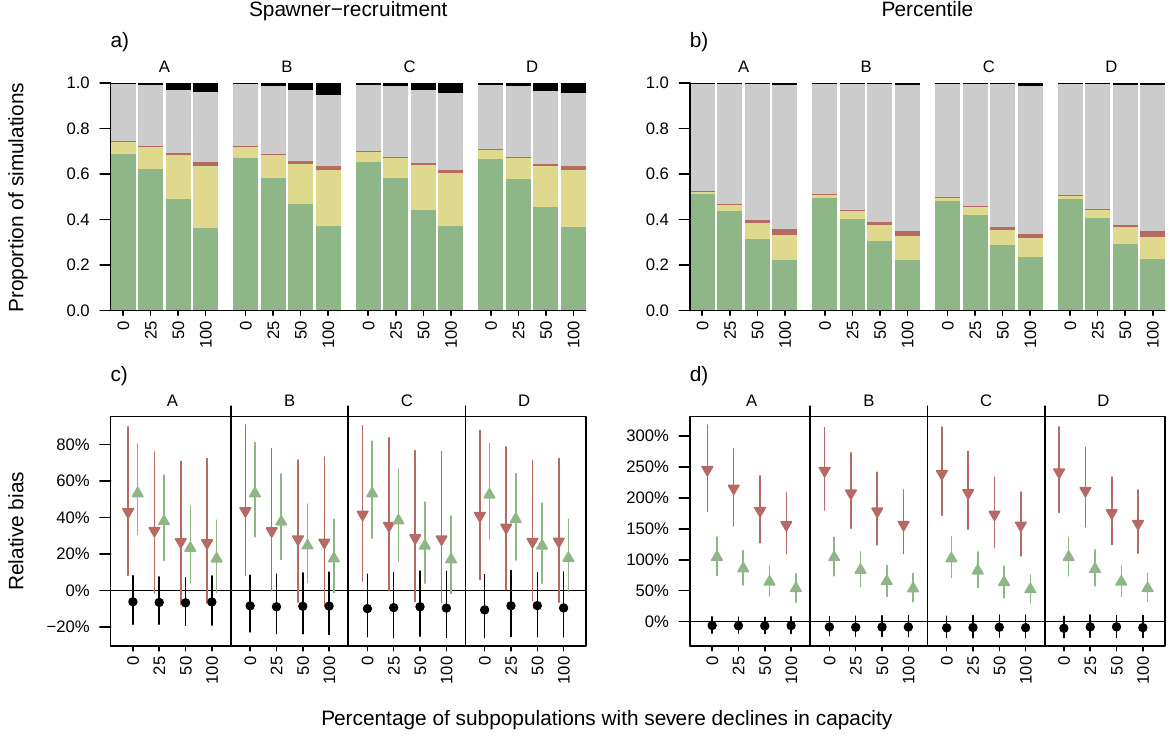

Figure S13. The effect of monitoring coverage (scenarios A-D; see below) and the percentage of populations with severe declines in capacity (bottom x-axis) on performance measures under the high productivity/HCR case. (a-b) The proportion of simulations with correct green, amber, or red status or biologically pessimistic misclassifications (grey) and biologically optimistic misclassifications (black) under the spawner-recruitment benchmarks (a) and percentile benchmarks (b). (c-d) The percent relative bias in the current spawner abundance (S_AVG_) and upper and lower benchmarks under the spawner-recruitment benchmarks (c) and the percentile benchmarks (d). Monitoring scenario A had no decline in coverage, B captured the observed decline in coverage of non-indicator streams, C had moderate declines in coverage of both indicator and non-indicator as observed for chum salmon on the north and central coast, and D had recent decline in coverage of indicator streams (Figure S6). These results for Scenarios A-B are also presented in the main text, Figure 8.

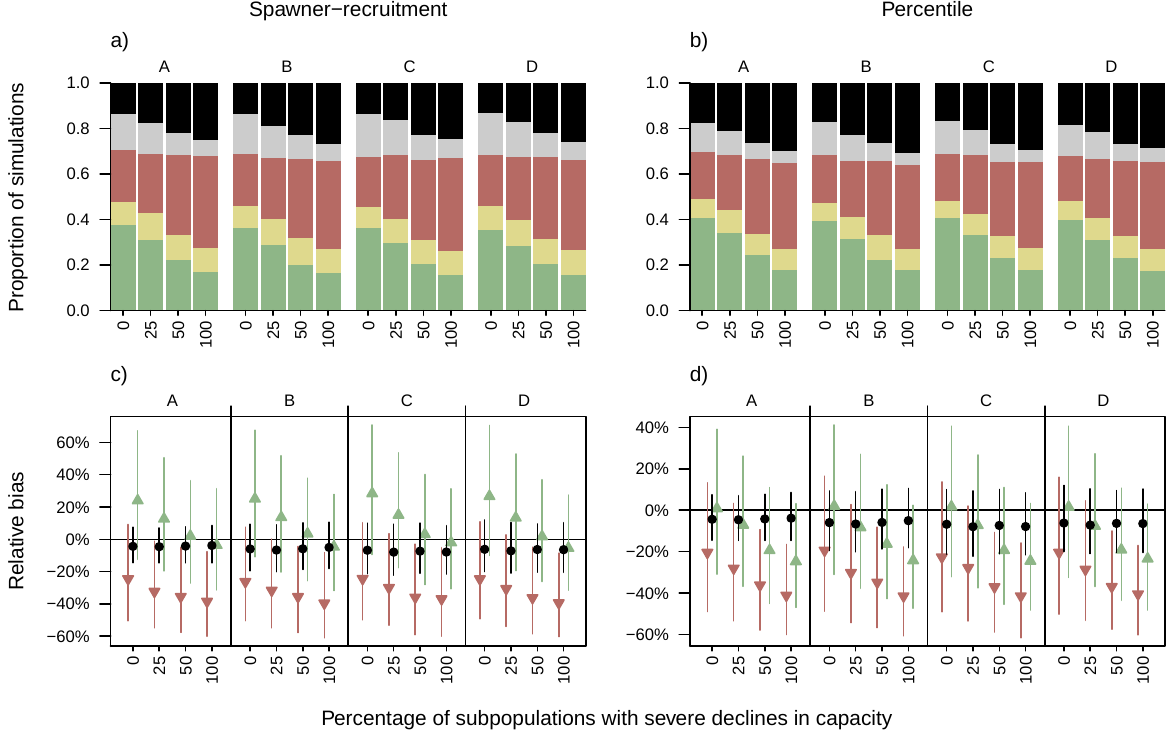

Figure S14.The effect of monitoring-coverage scenario (A-D; top x-axis) and the percentage of populations with severe declines in capacity (bottom x-axis) on performance measures **under the low-productivity moderate-harvest case**. (a-b) The proportion of simulations with correct green, amber, or red status or pessimistic misclassifications (grey) and optimistic misclassifications (black) under the spawner-recruitment benchmarks (a) and percentile benchmarks (b). (c-d) The percent relative bias in the current spawner abundance (S_AVG_) and upper and lower benchmarks under the spawner-recruitment benchmarks (c) and the percentile benchmarks (d). See Table 1 for a description of monitoring-coverage scenarios and declines in capacity.

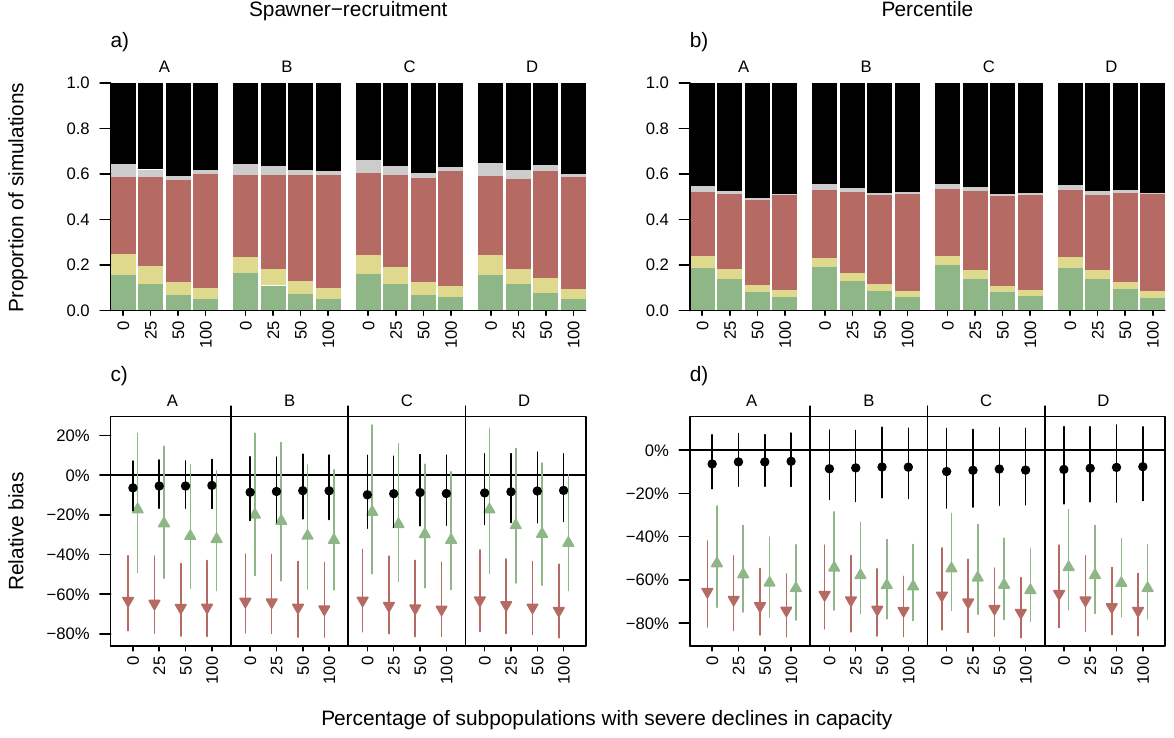

Figure S15. The effect of monitoring-coverage scenario (A-D; top x-axis) and the percentage of populations with severe declines in capacity (bottom x-axis) on performance measures **under the low-productivity high-harvest case**. (a-b) The proportion of simulations with correct green, amber, or red status or pessimistic misclassifications (grey) and optimistic misclassifications (black) under the spawner-recruitment benchmarks (a) and percentile benchmarks (b). (c-d) The percent relative bias in the current spawner abundance (S_AVG_) and upper and lower benchmarks under the spawner-recruitment benchmarks (c) and the percentile benchmarks (d). See Table 1 for a description of monitoring-coverage scenarios and declines in capacity.

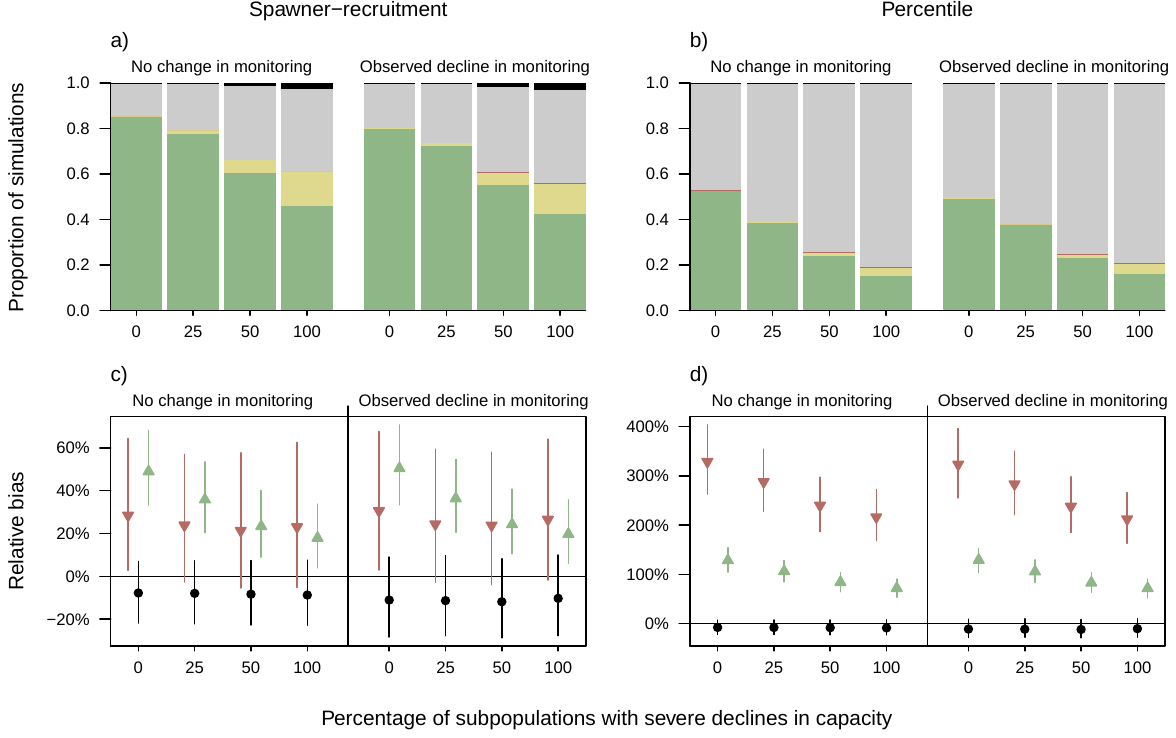

Figure S16. The effect of monitoring coverage (no change and decline; Table 2) and the percentage of spawning populations with severe declines in capacity (x-axis) on performance measures under the base case of high productivity and HCR (as in Fig. 8 of the main text**), assuming zero correlation in recruitment deviates among populations (**$\boldsymbol{\rho=0}$**; compared to default value of** $\boldsymbol{\rho=0.46}$ **in Fig. 8)**. (a-b) The proportion of simulations with correct green, amber, or red status or pessimistic misclassifications (grey) and optimistic misclassifications (black) under the spawner-recruitment benchmarks (a) and percentile benchmarks (b). (c-d) The percent relative bias (median $\pm$ interquartile range among 4000 MC simulations) in the current spawner abundance (S_AVG_; black) and lower and upper benchmarks (red and green, respectively) under the spawner-recruitment benchmarks (c) and the percentile benchmarks (d). See Online Supplement for results under the low-productivity high-harvest case.

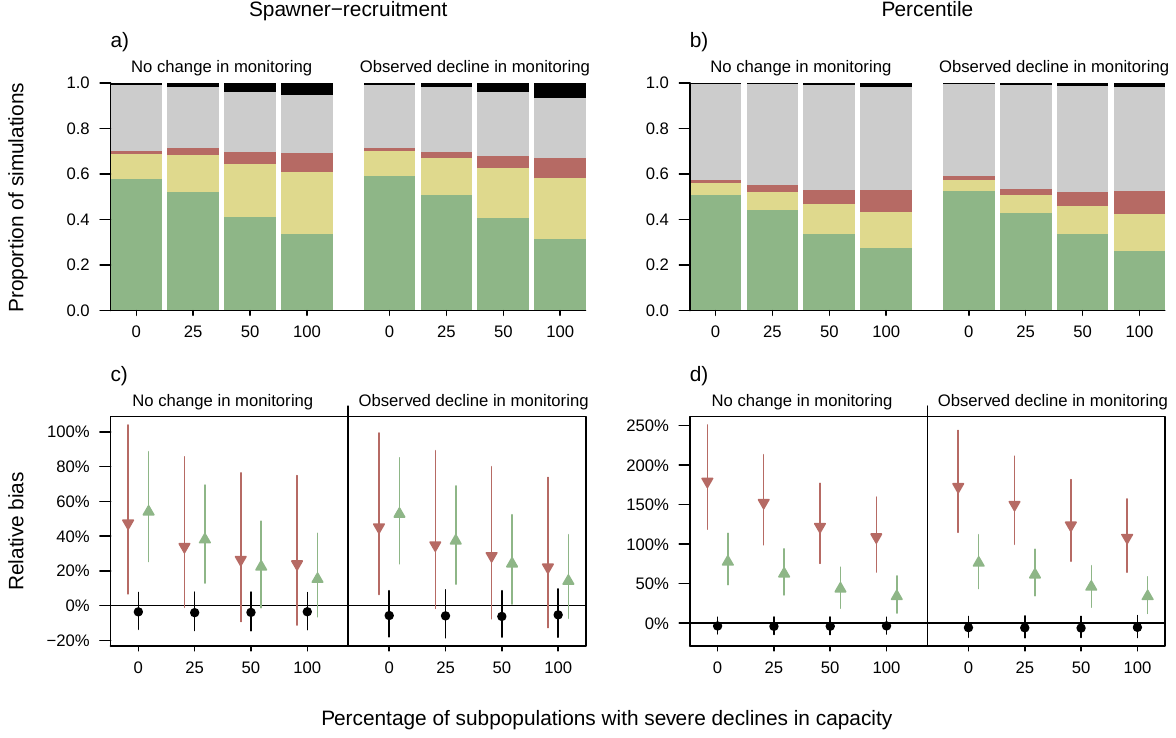

Figure S17. The effect of monitoring coverage (no change and decline; Table 2) and the percentage of spawning populations with severe declines in capacity (x-axis) on performance measures under the base case of high productivity and HCR (as in Fig. 8 of the main text**), assuming a high degree of correlation in recruitment deviates among populations (**$\boldsymbol{\rho=0.9}$**; compared to default value of** $\boldsymbol{\rho=0.46}$ **in Fig. 8)**. (a-b) The proportion of simulations with correct green, amber, or red status or pessimistic misclassifications (grey) and optimistic misclassifications (black) under the spawner-recruitment benchmarks (a) and percentile benchmarks (b). (c-d) The percent relative bias (median $\pm$ interquartile range among 4000 MC simulations) in the current spawner abundance (S_AVG_; black) and lower and upper benchmarks (red and green, respectively) under the spawner-recruitment benchmarks (c) and the percentile benchmarks (d). See Online Supplement for results under the low-productivity high-harvest case. Compare also to Figure S16, above, with $\rho=0$.

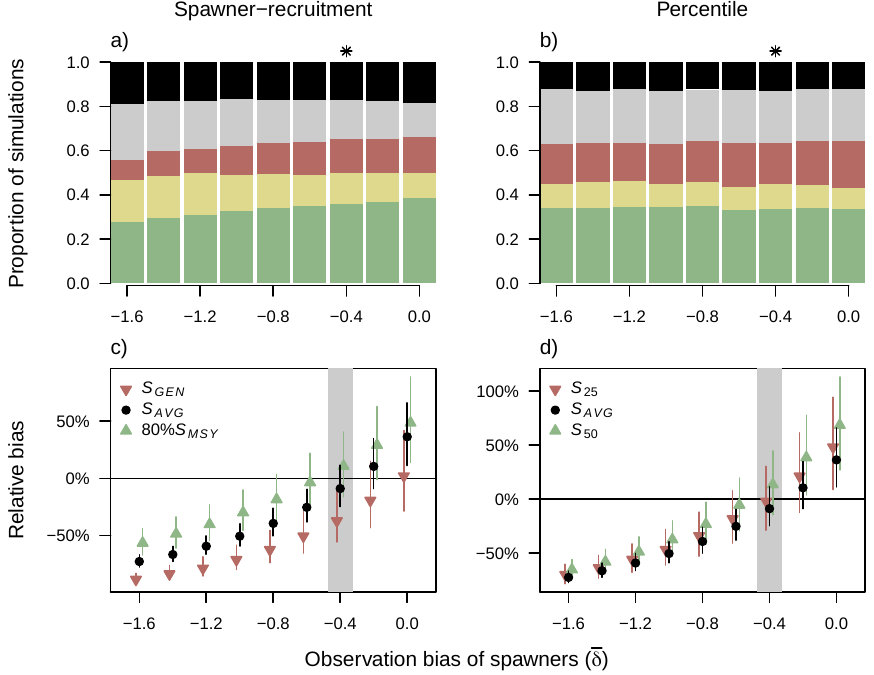

Figure S18. The effect of observation bias in the number of spawners (x-axis) on performance measures under the low-productivity moderate-harvest case. (a-b) The proportion of simulations with correct green, amber, or red status or pessimistic misclassifications (grey) and optimistic misclassifications (black) under the spawner-recruitment benchmarks (a) and percentile benchmarks (b). (c-d) The percent relative bias in the current spawner abundance (S_AVG_) and upper and lower benchmarks under the spawner-recruitment benchmarks (c) and the percentile benchmarks (d). The asterisk in (a-b) and grey zone in (c-d) indicate the default parameter value of $\bar{\delta}$ = -0.4, and the biasthat matches the Expansion Factor III of F’’’ = 1.5 applied in all simulations.

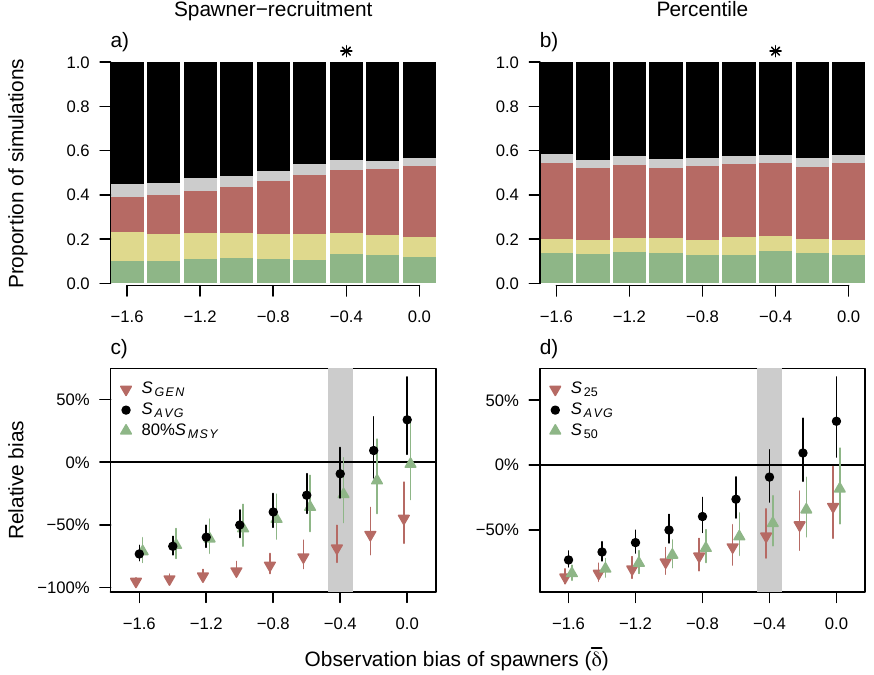

Figure S19. The effect of observation bias in the number of spawners (x-axis) on performance measures under the low-productivity high-harvest case. (a-b) The proportion of simulations with correct green, amber, or red status or pessimistic misclassifications (grey) and optimistic misclassifications (black) under the spawner-recruitment benchmarks (a) and percentile benchmarks (b). (c-d) The percent relative bias in the current spawner abundance (S_AVG_) and upper and lower benchmarks under the spawner-recruitment benchmarks (c) and the percentile benchmarks (d). The asterisk in (a-b) and grey zone in (c-d) indicate the default parameter value of $\bar{\delta}$ = -0.4, and the biasthat matches the Expansion Factor III of F’’’ = 1.5 applied in all simulations.

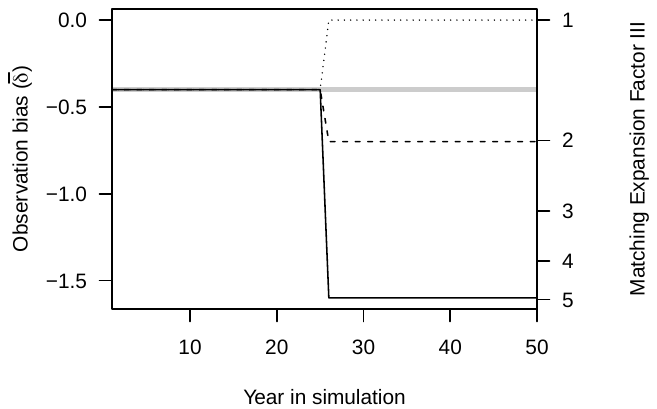

Figure S20. Illustration of four scenarios looking at a change in observation bias ($\bar{\delta}$, y-axis) halfway through simulations. Prior to the change, the observation bias matches the value of Expansion Factor III of F’’’ = 1.50 (right y-axis). After the change, the four scenarios consider: (1) a bias of $\bar{\delta}=$-1.6, corresponding to a value of F’’’ = 5.0 (solid black line), (2) $\bar{\delta}=$- 0.7, corresponding to a value of F’’’ = 2.0 (dashed black line), (3) no change (solid grey line), and (4) a decrease in bias to $\bar{\delta}=0$, corresponding to a value of F’’’ = 1.0 (dotted black line).

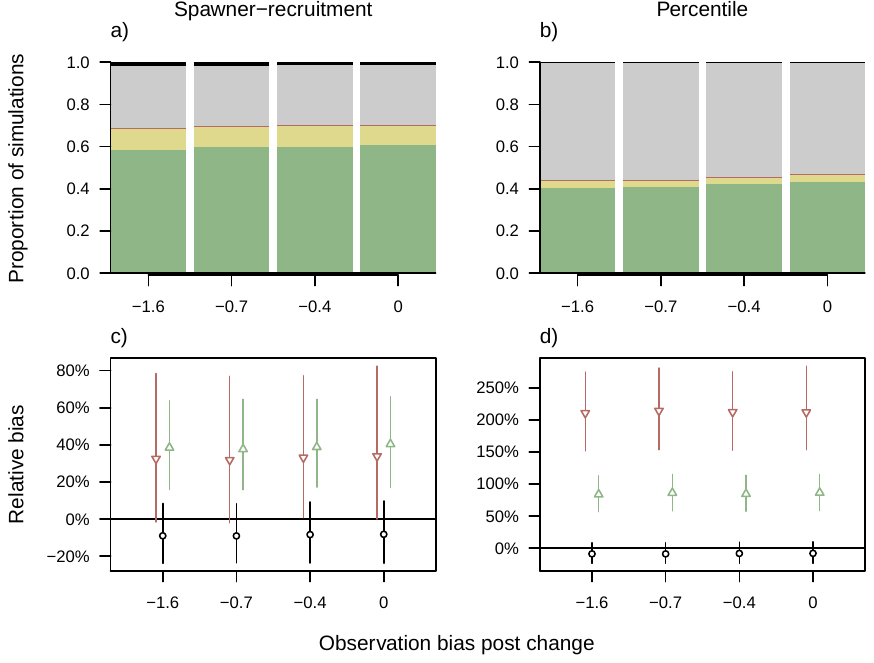

Figure S21. The effect of a change in observation bias of spawners halfway through the simulation (Figure S18) on performance measures under the high-productivity HCR case. (a-b) The proportion of simulations with correct green, amber, or red status or pessimistic misclassifications (grey) and optimistic misclassifications (black) under the spawner-recruitment benchmarks (a) and percentile benchmarks (b). (c-d) The percent relative bias in the current spawner abundance (S_AVG_) and upper and lower benchmarks under the spawner-recruitment benchmarks (c) and the percentile benchmarks (d). Results under other constant harvest rate cases were similar in that the change in observation bias had little effect on status outcomes.

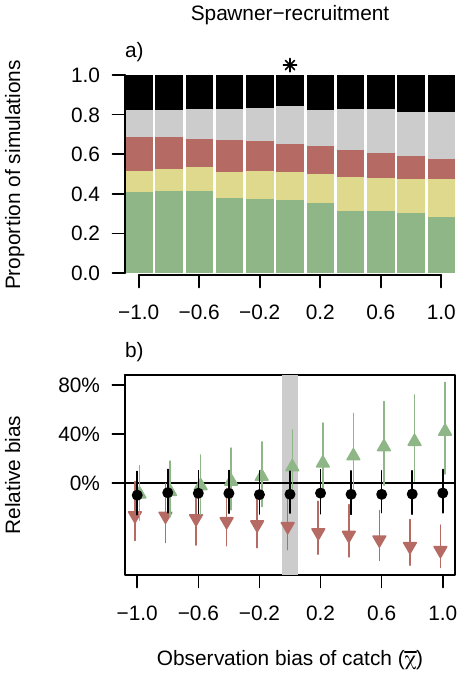

Figure S22. The effect of observation bias in catch (x-axis) on performance measures under the low productivity/moderate harvest. (a-b) The proportion of simulations with correct green, amber, or red status or pessimistic misclassifications (grey) and optimistic misclassifications (black) under the spawner-recruitment benchmarks (a) and percentile benchmarks (b). (c-d) The percent relative bias in the current spawner abundance (S_AVG_) and upper and lower benchmarks under the spawner-recruitment benchmarks (c) and the percentile benchmarks (d). The asterisk in (a-d) and grey zone in (c-d) indicate the default parameter value of $\bar{\chi}$ = 0.

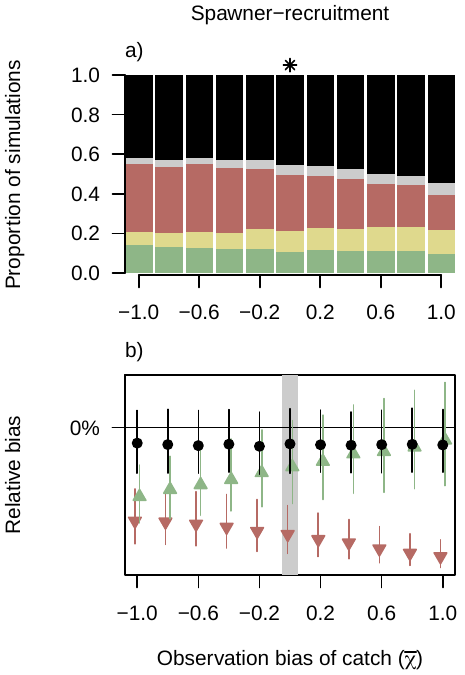

Figure S23. The effect of observation bias in catch (x-axis) on performance measures under the low-productivity high-harvest. (a-b) The proportion of simulations with correct green, amber, or red status or pessimistic misclassifications (grey) and optimistic misclassifications (black) under the spawner-recruitment benchmarks (a) and percentile benchmarks (b). (c-d) The percent relative bias in the current spawner abundance (S_AVG_) and upper and lower benchmarks under the spawner-recruitment benchmarks (c) and the percentile benchmarks (d). The asterisk in (a-d) and grey zone in (c-d) indicate the default parameter value of $\bar{\chi}$ = 0.

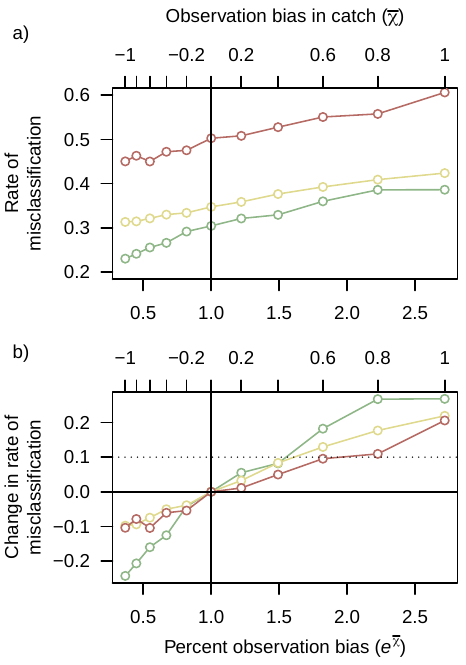

Figure S24. a) The rate of misclassification (e.g., sum of pessimistic and optimistic misclassifications from Figure S20 - Figure S21) over observation bias in catch ($\bar{\chi}$, top x-axis), also shown as the percent bias ($e^{\bar{\chi}}$, bottom x-axis) for three cases: base case of high productivity and HCR (green), low-productivity moderate-harvest (amber) and low-productivity high-harvest (red). Vertical solid black line indicates $\bar{\chi}=0$, or no bias in the observation of catch. b) The change in misclassifications from no catch bias over observation bias in catch ($\bar{\chi}$, top x-axis), also shown as the percent bias ($e^{\bar{\chi}}$, bottom x-axis) for the three cases. Horizontal solid black line indicates zero change in misclassifications for $\bar{\chi}=0$, and the dotted line indicates a 10% increase in misclassifications.

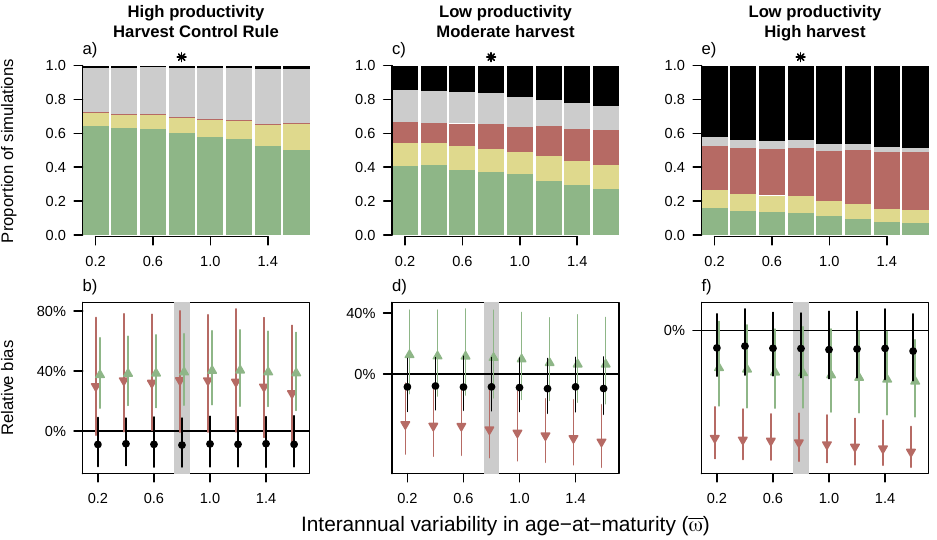

Figure S25. The effect of interannual variability in age-at-return (x-axis) on performance measures for the spawner-recruitment benchmarks under the three cases that we considered. (a-b) The proportion of simulations with correct green, amber, or red status or pessimistic misclassifications (grey) and optimistic misclassifications (black) under the spawner-recruitment benchmarks (a) and percentile benchmarks (b). (c-d) The percent relative bias in the current spawner abundance (S_AVG_) and upper and lower benchmarks under the spawner-recruitment benchmarks (c) and the percentile benchmarks (d). The asterisk in (a-d) and grey zone in (c-d) indicate the default parameter value of $\bar{\omega}$ = 0.8.

#### 3.1 Sensitivity to the variability in recruitment deviates in the true population dynamics

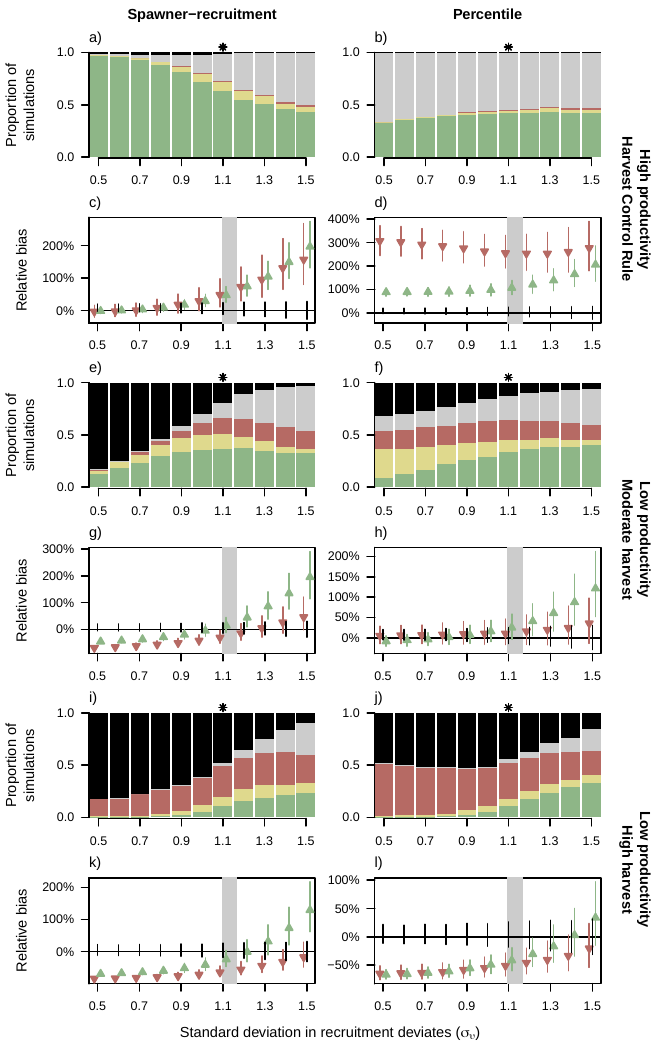

Figure S26. The effect of changes in the standard deviation in recruitment deviates (x-axis) on performance measures for the spawner-recruitment benchmarks (left) and percentile benchmarks (right) under the three cases that we considered. (high productivity/HCR, low productivity/moderate harvest, and low productivity/high harvest)). (a,b,e,f,I,j) The proportion of simulations with correct green, amber, or red status or pessimistic misclassifications (grey) and optimistic misclassifications (black). (c,d,g,h,k,l) The percent relative bias in the current spawner abundance (S_AVG_) and upper and lower benchmarks under the spawner-recruitment benchmarks (c) and the percentile benchmarks (d). The asterisk or grey zone indicate the default parameter value of $\sigma_{\upsilon}$ = 1.13.

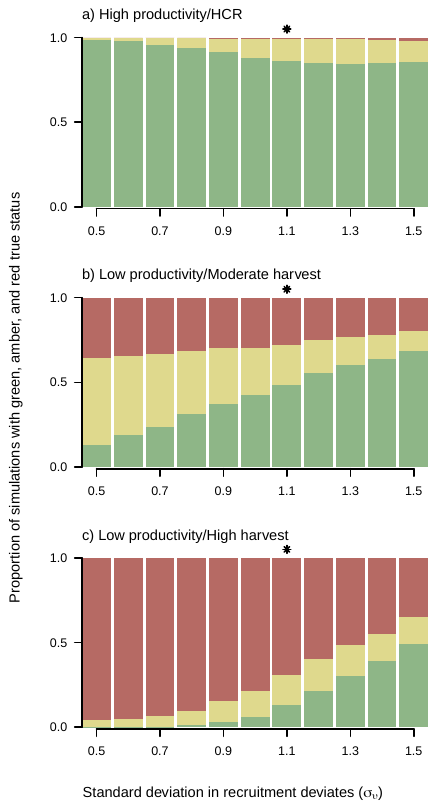

Figure S27. The proportion of simulations with a true status of green, amber, or red (y-axis) over increasing standard deviation in the recruitment deviates (x-axis) under the three cases (a-c). The asterisk indicates the default parameter value of $\sigma_{\upsilon}$ = 1.13.
